# Inferring population structure in biobank-scale genomic data

**DOI:** 10.1101/2021.05.11.443705

**Authors:** Alec M. Chiu, Erin K. Molloy, Zilong Tan, Ameet Talwalkar, Sriram Sankararaman

**Affiliations:** Bioinformatics Interdepartmental Program, University of California, Los Angeles, CA USA; Department of Computer Science, University of California, Los Angeles, CA USA; Institute for Advanced Computer Studies, University of Maryland, College Park, College Park, MD USA; Facebook, Inc., Menlo Park, CA USA; Machine Learning Department, Carnegie Mellon University, Pittsburgh, PA USA; Department of Human Genetics, University of California, Los Angeles, CA USA; Department of Computational Medicine, David Geffen School of Medicine, University of California, Los Angeles, CA USA

## Abstract

Inferring the structure of human populations from genetic variation data is a key task in population and medical genomic studies. While a number of methods for population structure inference have been proposed, current methods are impractical to run on biobank-scale genomic datasets containing millions of individuals and genetic variants. We introduce SCOPE, a method for population structure inference that is orders of magnitude faster than existing methods while achieving comparable accuracy. SCOPE infers population structure in about a day on a dataset containing one million individuals and variants as well as on the UK Biobank dataset containing 488,363 individuals and 569,346 variants. Furthermore, SCOPE can leverage allele frequencies from previous studies to improve the interpretability of population structure estimates.

## Introduction

Inference of population structure is a central problem in human genetics with applications ranging from fine-grained understanding of human history [1] to correcting for population stratification in genome-wide association studies (GWAS) [2]. Approaches to population structure inference [3, 4, 5, 6, 7, 8] typically formalize the problem as one of estimating admixture proportions of each individual and ancestral population allele frequencies given genetic variation data.

The growth of repositories of genetic variation data over large numbers of individuals has opened up the possibility of inferring population structure at increasingly finer resolution [9, 10]. For instance, the UK Biobank [9] contains genotype data from approximately half a million British individuals across millions of SNPs. This development has necessitated methods that can be applied to large-scale datasets with reasonable runtime and memory requirements. Existing methods, however, do not scale to these datasets. Thus, we have developed SCOPE (SCalable pOPulation structure inferencE) – a scalable method capable of inferring population structure on biobank-scale data.

SCOPE utilizes a previously proposed likelihood-free framework [8] that involves estimation of the individual allele frequency (IAF) matrix through a statistical technique known as latent subspace estimation (LSE) [11] followed by a decomposition of the estimated IAF matrix into ancestral allele frequencies and admixture proportions. SCOPE uses two ideas to substantially improve the scalability of this approach. First, SCOPE uses randomized eigendecomposition [12] to efficiently estimate the latent subspace. Specifically, SCOPE avoids the need to form matrices that are expensive to compute on or require substantial memory instead working directly with the input genotype matrix. Second, SCOPE leverages the insight that the resulting method involves repeated multiplications of the genotype matrix and uses the Mailman algorithm for fast multiplication of the genotype matrix[13].

We benchmarked the accuracy and efficiency of SCOPE on simulated and real datasets. In simulations, SCOPE obtains accuracy comparable to existing methods while being three to 1,800 times faster. Relative to the previous state-of-the-art scalable method (TeraStructure [7]), SCOPE is 3 to 144 times faster. SCOPE can estimate population structure in about a day for a dataset consisting of one million individuals and SNPs whereas TeraStructure, is extrapolated to require approximately 20 days. We used SCOPE to infer continental ancestry proportions (four ancestry groups) on the UK Biobank dataset (488,363 individuals and 569,346 SNPs) in about a day. We find that the inferred continental ancestry proportions are highly concordant with self-reported race and ethnicity (SIRE).

SCOPE additionally can be applied in a supervised setting. Given allele frequencies from reference populations [14, 15, 16], SCOPE can estimate admixture proportions corresponding to the reference populations, to enable greater interpretability.

## Results

### Accuracy

We assessed the accuracy of SCOPE using simulations under the Pritchard-Stephens-Donnelly (PSD) model [3] and a model of spatial structure [17]. We simulated several datasets using parameters calculated from two real datasets: the 1000 Genomes Project (TGP) [15] and the Human Genome Diversity Project (HGDP) [18] (benchmarking sections of Methods).

Under the PSD model, which matches the assumptions of the methods tested, ADMIXTURE is the most accurate followed by SCOPE and ALStructure (Figures 1, S1, S2, S3, S4). Among the scalable methods, TeraStructure and SCOPE, SCOPE tends to be more accurate in terms of both Jensen-Shannon divergence (JSD) (Table 1) and and root-mean-square error (RMSE) (Table 2). We also assessed accuracy under a spatial model, which violates the assumptions of the PSD model by inducing a spatial relationship between the admixture proportions (Figures 2, S5, S6, S7). Under this scenario, SCOPE and ALStructure are typically the most accurate (Tables 1, 2).

**Table 1:** Jensen-Shannon divergence measurements for methods on simulated data. Jensen-Shannon divergence (JSD) was computed against the ground truth admixture proportions for each simulation. Values are displayed as percentages rounded to one decimal place. Estimated proportions of 0 were set to 1 10^−9^ (see Methods). A ‘-’ denotes that the method was not run due to projected time or memory usage. Bold values denote the best value for each dataset.

| Dataset Type | Base Dataset | k | n | m | ADMIXTURE | fastStructure | TeraStructure | ALStructure | SCOPE |
| --- | --- | --- | --- | --- | --- | --- | --- | --- | --- |
| PSD | HGDP | 6 | 10,000 | 10,000 | **3.4** | 9.0 | 19.7 | 5.1 | 5.1 |
| PSD | TGP | 6 | 10,000 | 10,000 | **1.1** | 16.4 | 12.7 | 2.7 | 2.7 |
| PSD | TGP | 6 | 10,000 | 1,000,000 | **0.04** | 11.7 | 0.2 | - | 0.2 |
| PSD | TGP | 6 | 100,000 | 1,000,000 | - | - | 0.4 | - | **0.3** |
| PSD | TGP | 6 | 1,000,000 | 1,000,000 | - | - | - | - | **0.3** |
| Spatial | HGDP | 6 | 10,000 | 10,000 | 9.3 | 48.8 | 8.3 | **3.0** | 3.8 |
| Spatial | TGP | 6 | 10,000 | 10,000 | 9.9 | 44.9 | **3.5** | 4.9 | 4.7 |
| Spatial | TGP | 10 | 10,000 | 100,000 | 17.9 | 50.1 | 9.1 | 11.7 | **8.0** |
| Spatial | TGP | 10 | 10,000 | 1,000,000 | - | - | **9.5** | - | 10.4 |

**Table 2:** Root-mean-square error measurements for methods on simulated data. Root-mean-square error (RMSE) was computed against the ground truth admixture proportions for each simulation. RMSE is displayed in percentage and rounded to the first decimal place. A ‘-’ denotes that the method was not run due to projected time or memory usage. Bold values denote the best value for each dataset.

| Dataset Type | Base Dataset | k | n | m | ADMIXTURE | fastStructure | TeraStructure | ALStructure | SCOPE |
| --- | --- | --- | --- | --- | --- | --- | --- | --- | --- |
| PSD | HGDP | 6 | 10,000 | 10,000 | **4.0** | 10.3 | 16.6 | 5.6 | 5.6 |
| PSD | TGP | 6 | 10,000 | 10,000 | **1.8** | 15.9 | 13.7 | 3.2 | 3.2 |
| PSD | TGP | 6 | 10,000 | 1,000,000 | **0.2** | 12.4 | 0.9 | - | 0.3 |
| PSD | TGP | 6 | 100,000 | 1,000,000 | - | - | 1.0 | - | **0.4** |
| PSD | TGP | 6 | 1,000,000 | 1,000,000 | - | - | - | - | **0.5** |
| Spatial | HGDP | 6 | 10,000 | 10,000 | 11.9 | 31.1 | 10.2 | **5.7** | 6.5 |
| Spatial | TGP | 6 | 10,000 | 10,000 | 12.5 | 29.1 | **6.8** | 7.5 | 7.3 |
| Spatial | TGP | 10 | 10,000 | 100,000 | 10.8 | 22.8 | 8.8 | 8.5 | **6.7** |
| Spatial | TGP | 10 | 10,000 | 1,000,000 | - | - | 10.0 | - | **8.2** |

**Figure 1:**
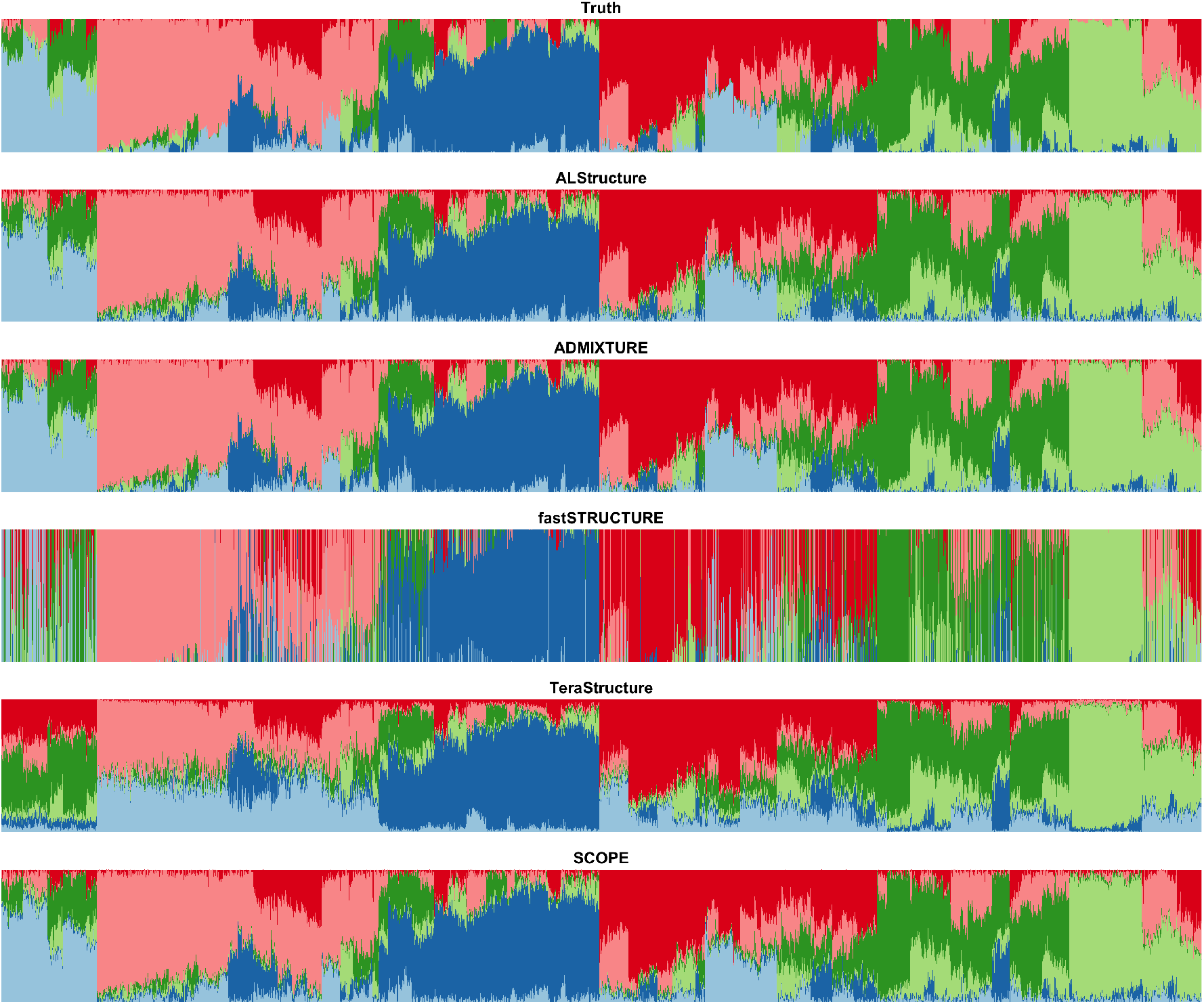
Population structure inference for simulations under PSD model generated using 1000 Genomes Phase 3 data. PSD model parameters were drawn from TGP data to generate a simulation dataset with 10,000 samples and 10,000 SNPs. The true admixture proportions and resulting inferred admixture proportions from each method are shown. Colors and order of samples are matched between each method to the truth.

**Figure 2:**
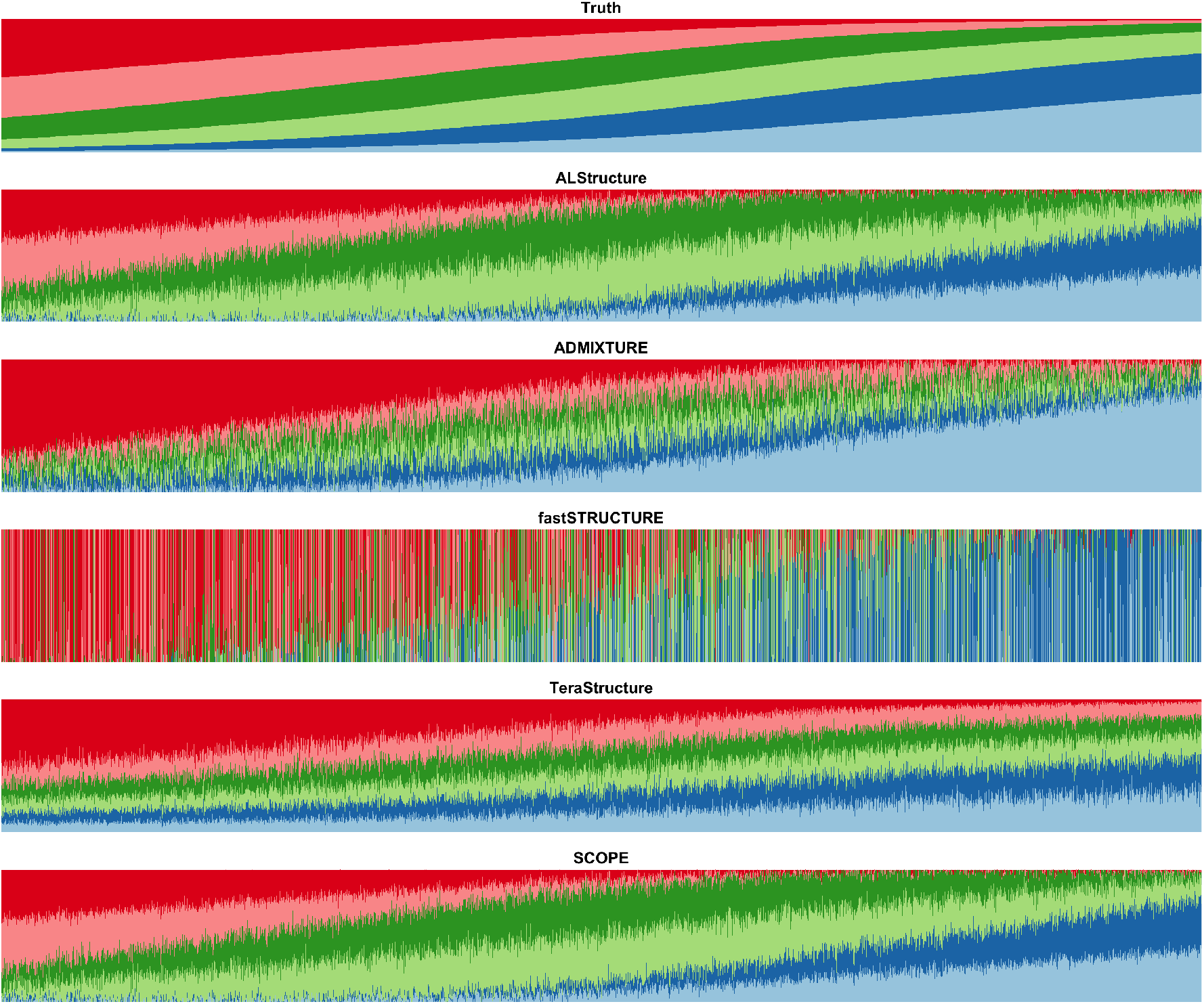
Population structure inference for simulations under a spatial model generated using 1000 Genomes Phase 3 data. Model parameters were drawn from TGP data to generate a simulation dataset with 10,000 samples and 10,000 SNPs under a spatial model (see Methods). The true admixture proportions and resulting inferred admixture proportions from each method are shown. Colors and order of samples are matched between each method to the truth.

### Runtime and memory

Using simulated and real datasets, we compared the runtime of SCOPE to ADMIXTURE, fast-Structure, TeraStructure, and ALStructure (Table 3). ADMIXTURE and fastStructure were not able to run on several of the larger datasets within practical time constraints while ALStructure could not be run on larger datasets due to memory constraints. On the largest PSD datasets that each method could be run on, SCOPE is over 150 times faster than ADMIXTURE (10,000 individuals by 1 million SNPs), over 500 times faster than fastStructure (10,000 individuals by 1 million SNPs), about 100 times faster than ALStructure (10,000 individuals by 100,000 SNPs), and over 110 times faster than TeraStructure (100,000 individuals by 1 million SNPs). SCOPE is also capable of running on a dataset containing one million SNPs and individuals in just over 24 hours (≈ 1 day) whereas TeraStructure is extrapolated to require about 500 hours (≈ 20 days) based on times reported in its manuscript [7] as well as our experiments (see benchmarking sections of Methods).

**Table 3:** Runtimes and fold-speedups of methods on simulations and real datasets. ADMIXTURE, TeraStructure, and SCOPE were run using 8 threads. ALStructure and fastStructure were run on a single thread due to their lack of multithreading implementations. Default parameters were used. TeraStructure's ‘-rfreq’ parameter was set to 10% of the number of SNPs. Times are rounded to the nearest minute and displayed in hours:minutes. The fold-speedup (runtime of method in seconds divided by runtime of SCOPE in seconds) achieved by SCOPE is denoted beneath each time in parentheses and rounded to the nearest integer. Bold values denote the best value for each dataset. Runtimes for SCOPE under one minute are denoted as “< 1 min.” A ‘-’ denotes that the method was not run due to projected time or memory usage.

| Dataset Type | Base Dataset | k | n | m | ADMIXTURE | fastStructure | TeraStructure | ALStructure | SCOPE |
| --- | --- | --- | --- | --- | --- | --- | --- | --- | --- |
| PSD | HGDP | 6 | 10,000 | 10,000 | 0:14<br>(48) | 3:44<br>(746) | 0:11<br>(36) | 0:30<br>(101) | < 1 min |
| PSD | TGP | 6 | 10,000 | 10,000 | 0:17<br>(206) | 1:22<br>(987) | 0:12<br>(144) | 0:23<br>(271) | < 1 min |
| PSD | TGP | 6 | 10,000 | 1,000,000 | 35:12<br>(156) | 114:51<br>(509) | 20:31<br>(91) | - | **0:14** |
| PSD | TGP | 6 | 100,000 | 1,000,000 | - | - | 237:02<br>(113) | - | **2:06** |
| PSD | TGP | 6 | 1,000,000 | 1,000,000 | - | - | - | - | **24:37** |
| Spatial | HGDP | 6 | 10,000 | 10,000 | 5:52<br>(440) | 4:06<br>(308) | 0:03<br>(3) | 1:39<br>(124) | < 1 min |
| Spatial | TGP | 6 | 10,000 | 10,000 | 3:11<br>(239) | 3:19<br>(249) | 0:07<br>(9) | 1:55<br>(144) | < 1 min |
| Spatial | TGP | 10 | 10,000 | 100,000 | 284:47<br>(1808) | 33:03<br>(210) | 4:29<br>(28) | 24:51<br>(158) | **0:09** |
| Spatial | TGP | 10 | 10,000 | 1,000,000 | - | - | 15:22<br>(9) | - | **1:47** |
| Real | HGDP | 10 | 940 | 642,951 | 4:24<br>(31) | 4:39<br>(33) | 0:40<br>(5) | 0:55<br>(7) | **0:08** |
| Real | HO | 14 | 1,931 | 385,089 | 13:28<br>(122) | 24:49<br>(224) | 1:37<br>(15) | 2:11<br>(20) | **0:07** |
| Real | TGP | 8 | 1,718 | 1,854,622 | 31:33<br>(33) | 8:53<br>(9) | 4:20<br>(5) | 11:16<br>(12) | **0:57** |
| Real | UKB | 4 | 488,363 | 569,346 | - | - | - | - | **25:57** |

The runtime of all methods increases under the spatial model. In this scenario, SCOPE is over 1800 times faster than ADMIXTURE (10,000 individuals by 100,000 SNPs), about 210 times faster than fastStructure (10,000 individuals by 100,000 SNPs), over 155 times faster than ALStructure (10,000 individuals by 100,000 SNPs), and about 9 times faster than TeraStructure (10,000 individuals by 1 million SNPs) on the largest dataset each method could be run on. Over all of the datasets, SCOPE is three to 1800 times faster than existing methods and three to 144 times faster than TeraStructure.

SCOPE has a reasonable memory footprint: for large datasets for which only TeraStructure and SCOPE were feasible, SCOPE uses slightly less memory than TeraStructure with the memory usage of SCOPE scaling linearly in the size of genotype matrix (*i.e.* the number of individuals times the number of SNPs) (Table S1). SCOPE requires less than 250 GB for the UK Biobank and 750 GB for the dataset consisting of one million individuals and SNPs.

### Accuracy of supervised analysis

Out of the methods tested, only SCOPE and ADMIXTURE are able to use supplied allele frequencies to perform population structure inference in a supervised fashion (Tables 4, S2). In the PSD model simulations, we observe a small improvement to both RMSE and JSD relative to unsupervised population structure inference (Figures S8, S9, S10, S11, S12). Under the spatial model simulations, the use of supervision obtains much greater accuracy compared to unsupervised inference (Figures 3, S13, S14, S15).

**Table 4:**
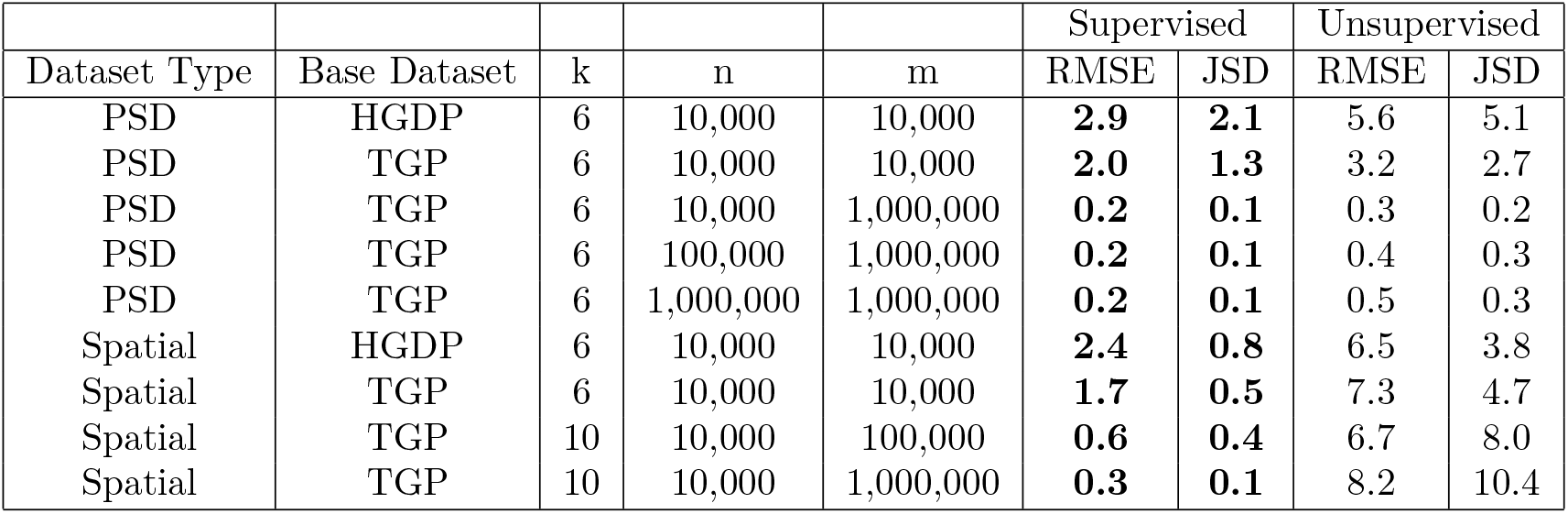
Accuracy of supervised population structure inference using supplied allele frequencies on simulations. True allele frequencies were supplied to SCOPE to use in supervised population structure inference. Root-mean-square error (RMSE) and Jensen-Shannon Divergence (JSD) were computed against the true admixture proportions. Estimated proportions of 0 were set to 1 10^−9^ for JSD calculations (see Methods). Values are displayed in percentages and rounded to the first decimal place. Bold values denote the best value for each dataset.

| Dataset Type | Base Dataset | k | n | m | Supervised |  | Unsupervised |  |
| --- | --- | --- | --- | --- | --- | --- | --- | --- |
|  |  |  |  |  | RMSE | JSD | RMSE | JSD |
| PSD | HGDP | 6 | 10,000 | 10,000 | **2.9** | **2.1** | 5.6 | 5.1 |
| PSD | TGP | 6 | 10,000 | 10,000 | **2.0** | **1.3** | 3.2 | 2.7 |
| PSD | TGP | 6 | 10,000 | 1,000,000 | **0.2** | **0.1** | 0.3 | 0.2 |
| PSD | TGP | 6 | 100,000 | 1,000,000 | **0.2** | **0.1** | 0.4 | 0.3 |
| PSD | TGP | 6 | 1,000,000 | 1,000,000 | **0.2** | **0.1** | 0.5 | 0.3 |
| Spatial | HGDP | 6 | 10,000 | 10,000 | **2.4** | **0.8** | 6.5 | 3.8 |
| Spatial | TGP | 6 | 10,000 | 10,000 | **1.7** | **0.5** | 7.3 | 4.7 |
| Spatial | TGP | 10 | 10,000 | 100,000 | **0.6** | **0.4** | 6.7 | 8.0 |
| Spatial | TGP | 10 | 10,000 | 1,000,000 | **0.3** | **0.1** | 8.2 | 10.4 |

**Figure 3:**
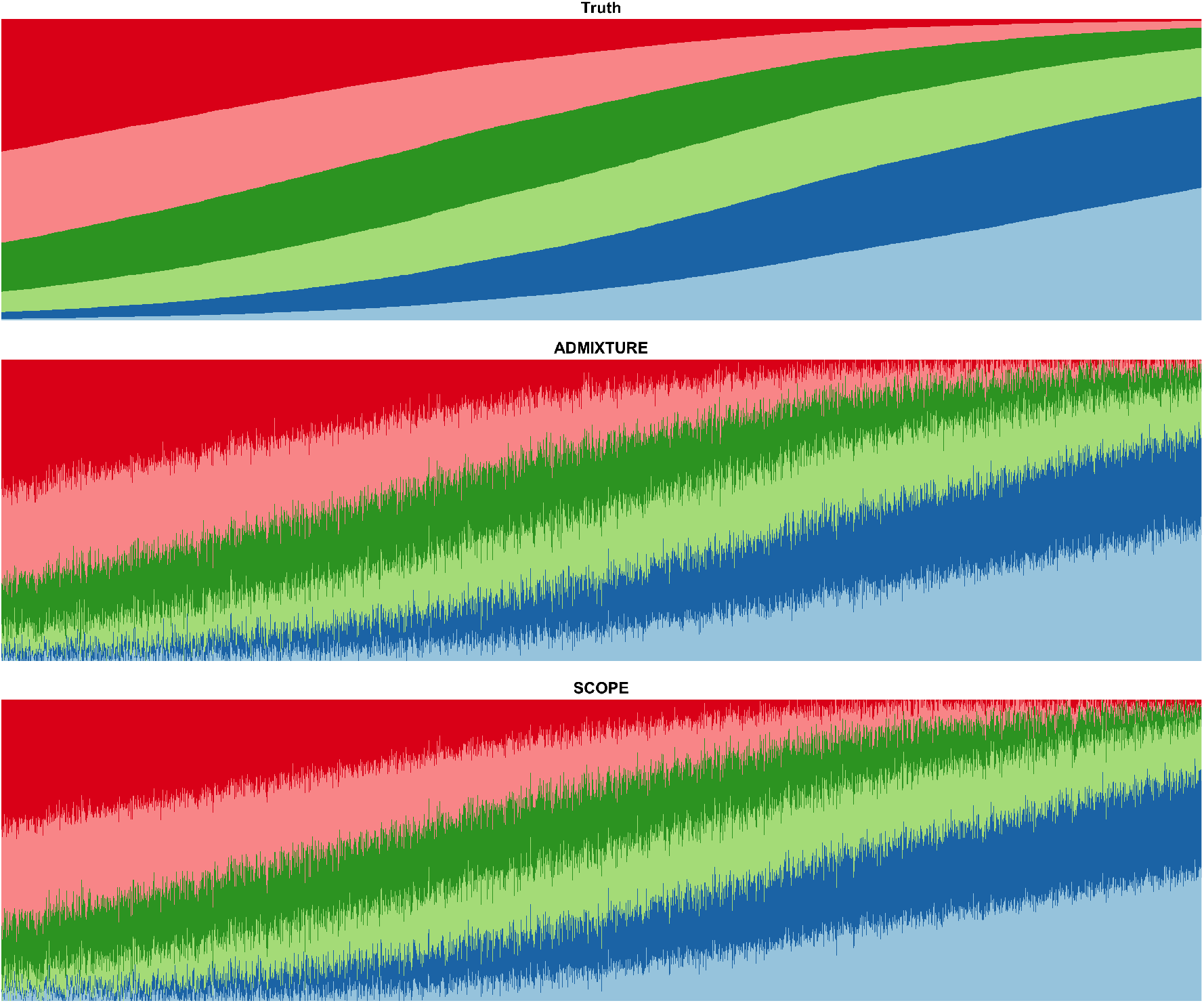
Supervised population structure inference for simulations under a spatial model generated using 1000 Genomes Phase 3 data. Model parameters were drawn from TGP data to generate a simulation dataset with 10,000 samples and 10,000 SNPs under a spatial model. Both methods were provided the true population allele frequencies as input. Colors and order of samples are matched between each method to the truth.

### Application to real genotype data

We applied SCOPE to several real, genomic datasets: TGP with 8 latent populations (*k* = 8) (Figure S16), HGDP with 10 populations (*k* = 10) (Figure S17), Human Origins (HO) [19] with 14 populations (*k* = 14) (Figure S18), and the UK Biobank with 4 populations (*k* = 4) (Figure 4) (see Methods). We chose the number of latent populations to be consistent with previous studies on these datasets [8, 7]. For the UK Biobank analysis, we chose four latent populations to infer continental ancestry groups. In terms of runtime and memory, we continued to observe trends consistent with our simulations where SCOPE is orders of magnitude faster than other methods while consuming reasonable amounts of memory (Tables 3, S1). We note that the runtime for inference on the UK Biobank is about the same as the runtime for our 1 million individual and SNP simulation despite the UK Biobank being approximately a quarter of its size, consistent with the increase in runtimes with model deviations as seen in the context of spatial simulations.

**Figure 4:**
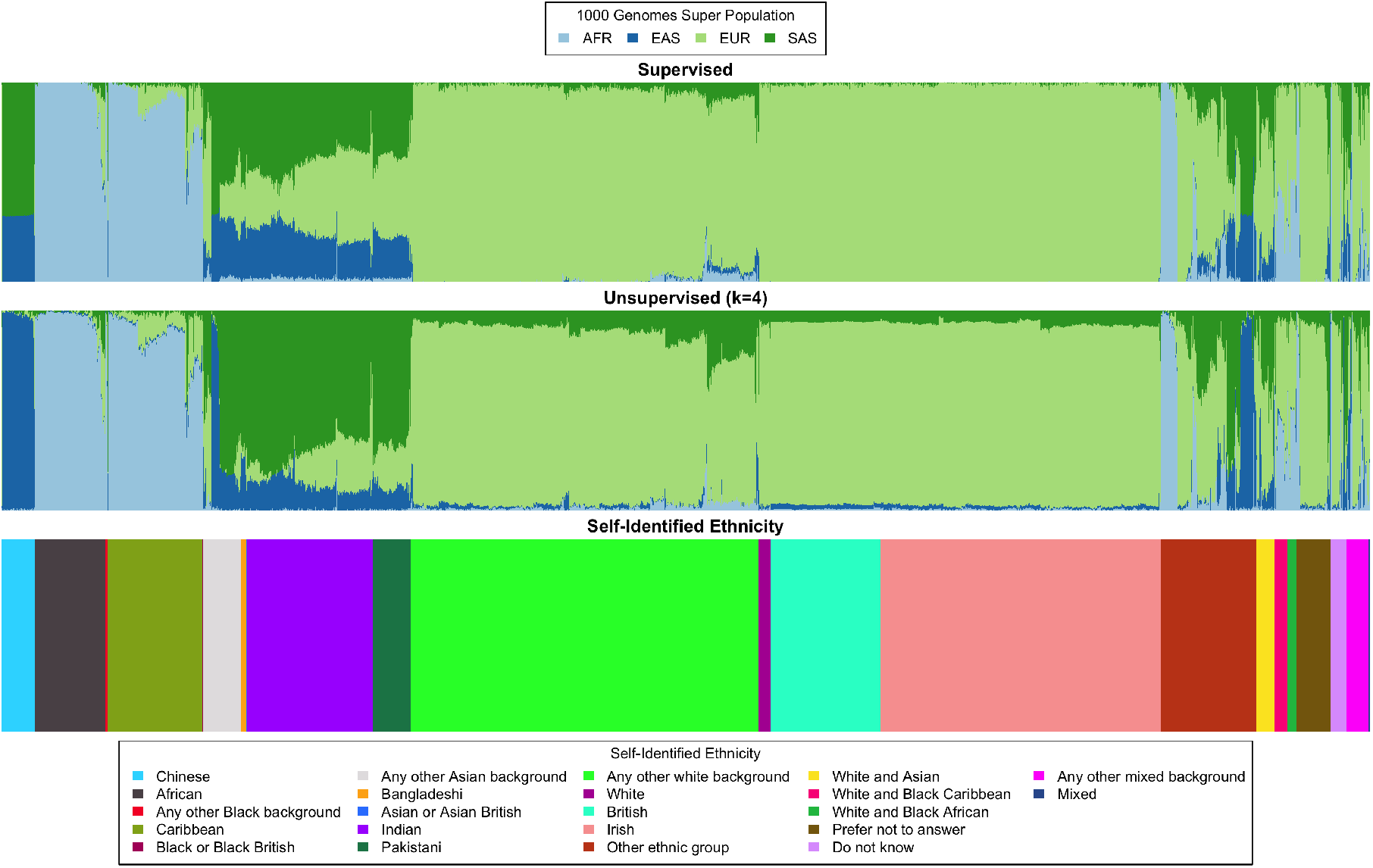
Continental ancestry inference on the UK Biobank. We ran population structure inference using SCOPE both supervised using 1000 Genomes Phase 3 allele frequencies (top) and unsupervised with 4 latent populations (middle). For reference, we plot the self-identified race/ethnicity (bottom). For visualization purposes, we reduced the number of self-identified British individuals to a random subset of 5, 000 individuals. Colors and order of samples are matched between each row of the figure. The full figure without individuals removed can be found in Figure S19.

Since there is no ground truth to assess accuracy on these datasets, we used concordance between SIRE and inferred admixture proportions as a metric. We trained multinomial logistic regression models to predict continental ancestry for the TGP (5 populations) and HGDP (7 populations) using the inferred admixture proportions from each method (Table S3). We find that all methods perform similarly on both datasets. For the UK Biobank, SCOPE is able to obtain 88.27% accuracy when using labels provided by UK Biobank (22 labels) and 95.75% accuracy when ambiguous/heterogeneous labels (*e.g.* Other, Mixed) are removed and population labels are collapsed to continental groupings (8 labels). We did not perform this analysis for the HO dataset due to several population labels only containing one sample.

We also utilized the supervised mode of SCOPE using known population allele frequencies from TGP superpopulations to infer continental ancestry for all individuals in the UK Biobank. We find that the supervised mode of SCOPE largely agreed with the unsupervised inference (Figures 4, S19).

## Discussion

We have presented SCOPE, a scalable method for inferring population structure from biobank-scale genomic data. We show that SCOPE remains accurate while being scalable in terms of runtime and memory requirements. SCOPE is also able to perform supervised analyses that leverage allele frequency estimates from previous studies to improve interpretability, runtime, and accuracy.

The methodology used in SCOPE can also be extended in several ways. Several methods that perform structure inference on other genomic datasets [20, 21] utilize semi-supervised approaches where there are both known and unknown populations. A possible approach for semi-supervision using SCOPE is to perform a multi-stage inference procedure where supervised inference is first applied and unsupervised inference is applied on the residual or unexplained structure. Most current methods, including SCOPE, ignore additional information within the data such as correlation patterns (*i.e.* linkage disequilibrium or LD). Some methods such as fineSTRUCTURE [22] can perform linkage-disequilibrium aware population structure inference but are challenging to scale. Methods that can model LD while retaining scalability is a key step in advancing population structure inference. Finally, extensions of the techniques used in SCOPE can be used to infer relevant structure in other domains such as metagenomics and single-cell transcriptomics.

## Code Availability

SCOPE can be found at https://github.com/sriramlab/SCOPE. Scripts for simulations, visualization, assessment, real data filtering, and additional code used in this study can be found at the repository as well.

## Methods

### The Structure/Admixture Model

The structure/admixture model links the *m* × *n* genotype matrix ***X*** (where rows refer to single nucleotide polymorphisms (SNPs) and columns refer to individual diploid genotypes, *x_ij_* ∈ {0, 1, 2}, *i* ∈ {1, …, *m*}, *j* ∈ {1, …, *n*}) to the *m* × *n* individual allele frequency (IAF) matrix ***F***, *m* × *k* ancestral population allele frequencies ***P***, and the *k* × *n* individual admixture proportions ***Q***(also termed the global ancestry of an individual). Here *m* denotes the number of SNPs, *n* denotes the number of individuals, and *k* denotes the number of latent populations. The IAF matrix, ancestral allele frequencies, and admixture proportions are mathematically related as ***F = PQ***. Furthermore, there are constraints on ***P*** and ***Q***. Each element of ***P*** is constrained to lie between 0 and 1 (0 ≤ *p_il_* ≤ 1, *i* ∈ {1, …, *m*}, *l* ∈ {1, …, *k*}). Each element of ***Q*** is non-negative (*q_lj_* ≥ 0, *l* ∈ {1, …, *k*}, *j* ∈ {1, …, *n*}) and the admixture proportion of each individual must sum to one (Σ_*l*_ *q_lj_* = 1). Finally, each entry of the genotype matrix is an independent draw from the corresponding entry of the IAF matrix ***F*** as: *x_ij_*|*f_ij_* ~ Binomial(2, *f_ij_*). The goal of population structure inference under the structure/admixture model is to estimate ***P*** and ***Q*** given ***X***.

### SCOPE

For scalable inference, SCOPE uses as its starting point a likelihood-free estimator of population structure previously proposed in ALStructure [8]. This estimator has two major steps: latent subspace estimation (LSE) and alternating least squares (ALS). LSE attempts to estimate the subspace spanned by the rows of ***Q*** [11] by computing a low-rank approximation to the matrix 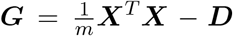 where each entry *d_j_* of the *n* × *n* diagonal matrix ***D*** is obtained as 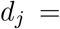 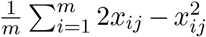. The latent subspace of ***Q*** is estimated as the span of the top *k* eigenvectors of ***G***: ***ν***_1_, …, ***ν****_k_*. After obtaining the top *k* eigenvectors ***V*** = [***ν***_1_, …, ***ν***_*k*_], ALStructure projects the data ***X*** onto ***ν*** to obtain an estimate of 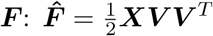. Truncated alternating least squares (ALS) is used to factorize the estimate, 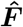, into estimates of ***P*** and 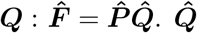 are the estimates of the individual admixture proportions.

A naive approach to compute the top *k* eigenvectors of ***G*** would involve first forming the matrix ***G*** and then computing its top *k* eigenvectors which would require 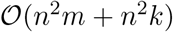 (if a full SVD is performed, this step would require 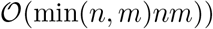. To perform scalable LSE, SCOPE uses techniques from randomized linear algebra [12], specifically the implicitly restarted Arnoldi method [23], to obtain the top *k* eigenvectors. This step involves repeatedly multiplying estimates of the eigenvectors ν*_l_* : *l* ∈ {1*, …, k*} with the genotype matrix: 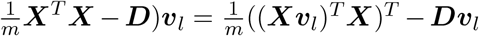 and can be performed without explicitly forming the matrix ***G***. Instead, this approach requires repeatedly computing 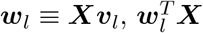, and ***Dν***_*l*_ which can be computed in 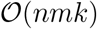 time. We use the C++ Spectra library (https://spectralib.org/) to implement these computations in SCOPE.

To efficiently compute 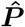 and 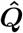 using truncated ALS, the matrix 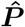 is initialized randomly with all values between 0 and 1 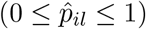. We iteratively solve for estimates of ***P*** and ***Q***, projecting the estimates onto the constraint space until convergence:

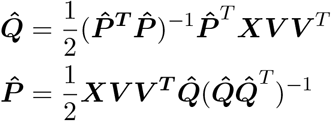

All values in 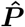 are truncated to be between 0 and 1 while 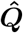 is projected onto the appropriate simplex. Each step of the ALS algorithm has runtime 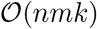.

Each of the computations in SCOPE require multiplying a genotype matrix with entries consisting of only 0, 1, and 2 for diploid genotype. These operations can be efficiently performed using the Mailman algorithm [13] that provides computational savings when there are repeated multiplications involving a matrix with a finite alphabet. We utilize the Mailman algorithm in computations involving the genotype matrix in both LSE and ALS so that the final time complexity of SCOPE is 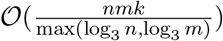.

### Supervised Population Structure Inference

SCOPE can utilize allele frequencies from reference populations to infer corresponding admixture proportions. In this scenario, we assume 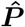, the population allele frequencies, are known. As a result, one only needs to compute 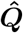 using the supplied 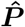. This allows the admixture proportions corresponding to the reference populations to be inferred in a single step of ALS once the LSE step is completed.

### Permutation Matching of Inferred Results

The output of population structure inference methods can result in output that is permuted even between different runs of the same method. It is critical to correctly match latent populations between methods and runs in order to properly assess results. To perform permutation matching, we employed a strategy similar to that of [24]. This permutation matching problem is better known as the Assignment Problem, which can be solved efficiently using linear programming. We first construct a score matrix using the distance metric created in [24]. The optimal permutation match can then be found by optimizing the total score from assignments through linear programming. We utilize the *lpSolve* (https://CRAN.R-project.org/package=lpSolve) package in R to solve the linear program.

### PSD Model Simulations

We perform simulations under the Structure or Pritchard-Stephens-Donnely (PSD) model [25]. In the PSD model, priors are placed on ***P*** and ***Q***:

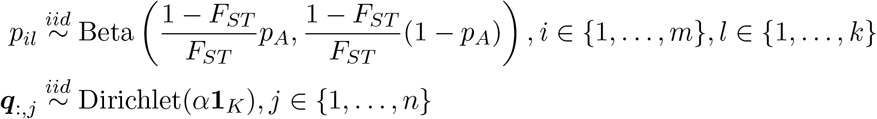

The allele frequencies *p_il_* are drawn from the Balding-Nichols model [26], which is a Beta distribution parametrized by the fixation index (*F_ST_*) and an initial allele frequency (*p_A_*). For our simulations, we calculated *F*_*ST*_ and *p_A_* from our real datasets. Admixture proportions ***q***:*,j* are drawn at random from a Dirichlet distribution. We take the product of the two matrices to form the IAF matrix, ***F*** = ***PQ***, and draw each genotype from a Binomial distribution parametrized by entries of ***F***: *x_ij_* ~ Binomial(2, *f_ij_*).

### Spatial Model Simulations

We also perform simulations under a spatial model similar to that in [17]. In the spatial model, allele frequencies *p_il_* are drawn as in the PSD model, but the admixture proportions, ***q***, are drawn from a 1D geography.

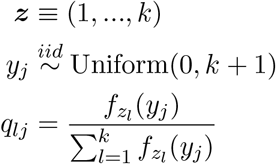

Populations are placed at integer values on a line. We get the resulting population position vector, ***z*** ≡ (1, …, *k*). Each individual has a position, *y_j_* drawn from a uniform distribution between 0 and *k* + 1. Proportions for each population are generated by using a normal distribution, where 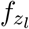 denotes the normal density function using *z_l_* (*l* ∈ {1, …, *k*}) as the mean and *σ*^2^ as variance. The resulting vector of proportions is then normalized to satisfy the constraints on ***Q***. We used *σ* = 4 for our simulations.

### Assessment of Results

We assess our results using two metrics: average Jensen-Shannon divergence (JSD) and average root-mean-square error (RMSE). We calculate the metrics between the true global ancestry proportions, ***Q*** and the estimates, 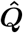, after 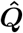 has been permutation matched to the true proportions.

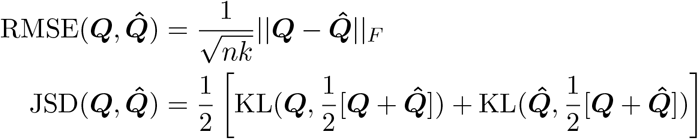

||·||*_F_* represents the Frobenius norm. KL is the Kullback-Leibler divergence, which is defined as:

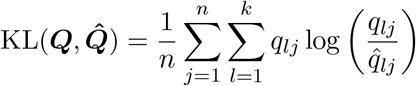

In the JSD calculations, we replace values of 0 in ***Q*** or 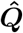 with 1 × 10^−9^ to avoid numerical issues.

### Datasets

We use the 1000 Genomes Project (TGP) [15, 14, 16], Human Origins (HO) [19], Human Genome Diversity Project (HGDP) [27, 28], and the UK Biobank (UKB) [9] in this study. The HGDP dataset was filtered to only include individuals in the H952 set [29], greater than 95% genotyping rate, and greater than 1% minor allele frequency (MAF). The TGP dataset was filtered to include unrelated individuals, greater than 95% genotyping rate, and greater than 1% MAF. The HO dataset was filtered for human-only samples, greater than 99% genotyping rate, and greater than 5% MAF. For the UK Biobank, we filtered for greater than 1% MAF, long range linkage disequilibrium (LD), and pairwise LD pruning in 50 kilobase windows, 80 variant step size, and an *r*^2^ threshold of 0.1. We calculate metrics such as *F_ST_* from the provided population and superpopulation labels provided by each dataset. To perform our supervised analyses, we use the common SNPs between the datasets involved. All genotype processing was performed using PLINK [30].

### Visualization of Results

We visualize our inferred admixture proportions as stacked bar plots. Estimates from all methods were permutation matched to enable easy comparison. For our PSD simulations, we performed hierarchical clustering with complete linkage on a Euclidean distance matrix calculated from the true admixture proportion matrix (***Q***) to obtain the order of samples. For our spatial simulations, we sorted by decreasing membership of the first population. For our real datasets, we perform the same hierarchical clustering strategy used for our PSD simulations, but use the estimates from ADMIXTURE 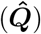 in place of the true admixture proportions. For the UK Biobank, we first took the average proportions for each SIRE group and performed hierarchical clustering on the averages to determine the order of the SIRE groups. We then performed hierarchical clustering within each SIRE group to determine the order of individuals within groups. For large datasets, we utilized *genieclust* [31], a scalable method for hierarchical clustering.

### Benchmarking

We compared SCOPE to ADMIXTURE v1.3.0 [5], fastSTRUCTURE [6], TeraStructure [7], and ALStructure v0.1.0 [8].

ADMIXTURE computes maximum-likelihood estimates while TeraStructure and fastSTRUCTURE compute approximate posterior estimates in a Bayesian model using variational inference. ALStructure, the framework which SCOPE builds upon, utilizes a two-stage strategy of first performing dimensionality reduction (latent subspace estimation) followed by matrix factorization (alternating least squares).

Each method was run with 8 threads with the exception of fastSTRUCTURE and ALStructure, which do not have multi-threaded implementations. Default parameters were used. TeraStructure has an additional ‘rfreq’ parameter, which was set to 10% of the number of SNPs as recommended by its authors. For SCOPE, we used convergence criteria of either 1,000 iterations of the ALS algorithm or a change between iterations less than 1 × 10^−5^, which we calculate as the RMSE between the estimated admixture matrices between two iterations. All experiments were performed on a server with two AMD EPYC 7501 32-Core Processors and 1 terabyte of RAM.

## Acknowledgments

We would like to thank Bogdan Pasaniuc and members of the Pasaniuc and Sankararaman labs for advice and comments on this project. This research was conducted using the UK Biobank Resource under application 33127. This work was funded by NIH grants T32HG002536 (A.M.C.), R35GM125055 (E.K.M., S.S.), an Alfred P. Sloan Research Fellowship (S.S.), and NSF grants DGE-1829071 (A.M.C.), III-1705121 (A.T. and S.S.).

