## Supplemental Information for "Inferring population structure in biobank-scale genomic data"

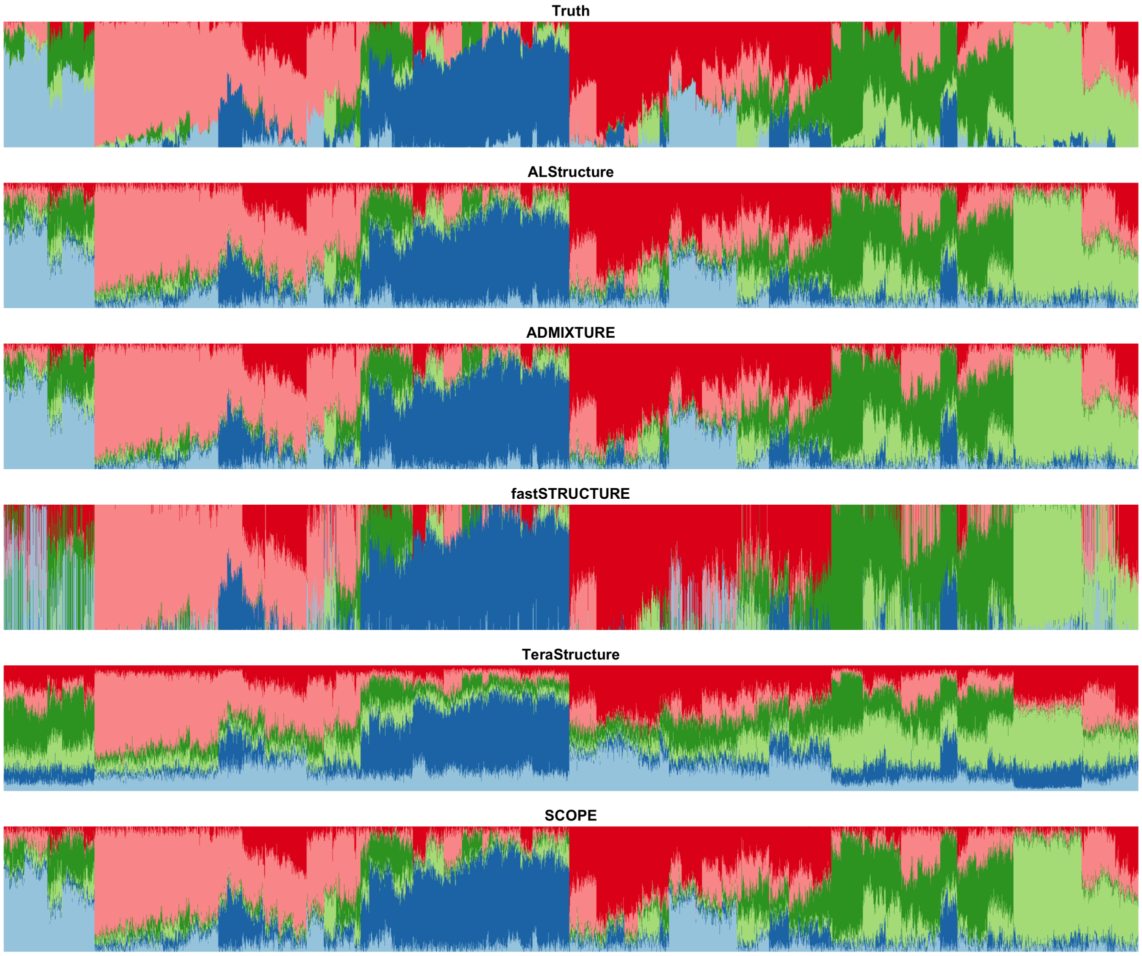


Figure S1: **Population structure inference for simulations under PSD model generated using Human Genomes Diversity Project data.** PSD model parameters were drawn from HGDP data to generate a simulation dataset with 10,000 samples and 10,000 SNPs. The true admixture proportions and resulting inferred admixture proportions from each method are shown. Colors and order of samples are matched between each method to the truth.

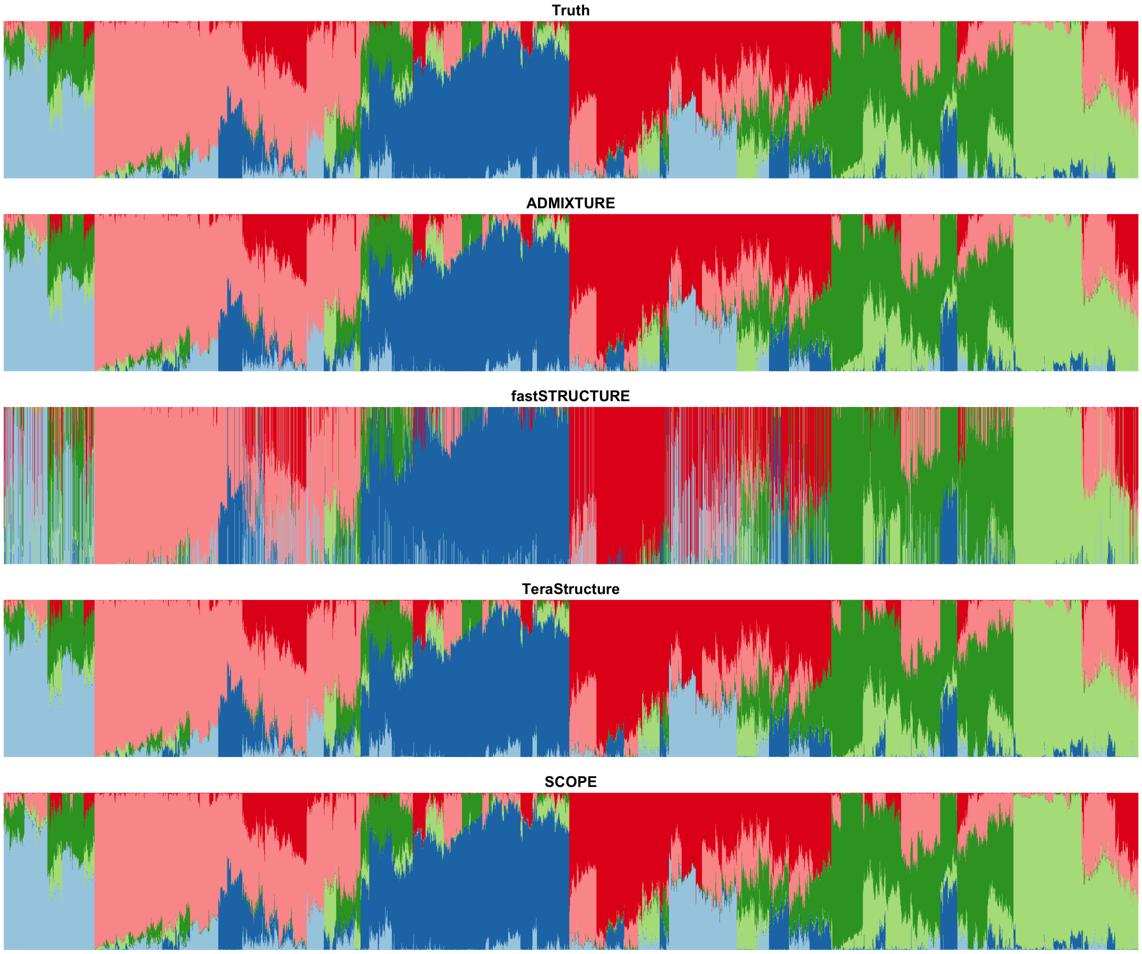


Figure S2: **Population structure inference for simulations under PSD model generated using 1000 Genomes Phase 3 data.** PSD model parameters were drawn from TGP data to generate a simulation dataset with 10,000 samples and 1 million SNPs. The true admixture proportions and resulting inferred admixture proportions from each method are shown. Colors and order of samples are matched between each method to the truth.

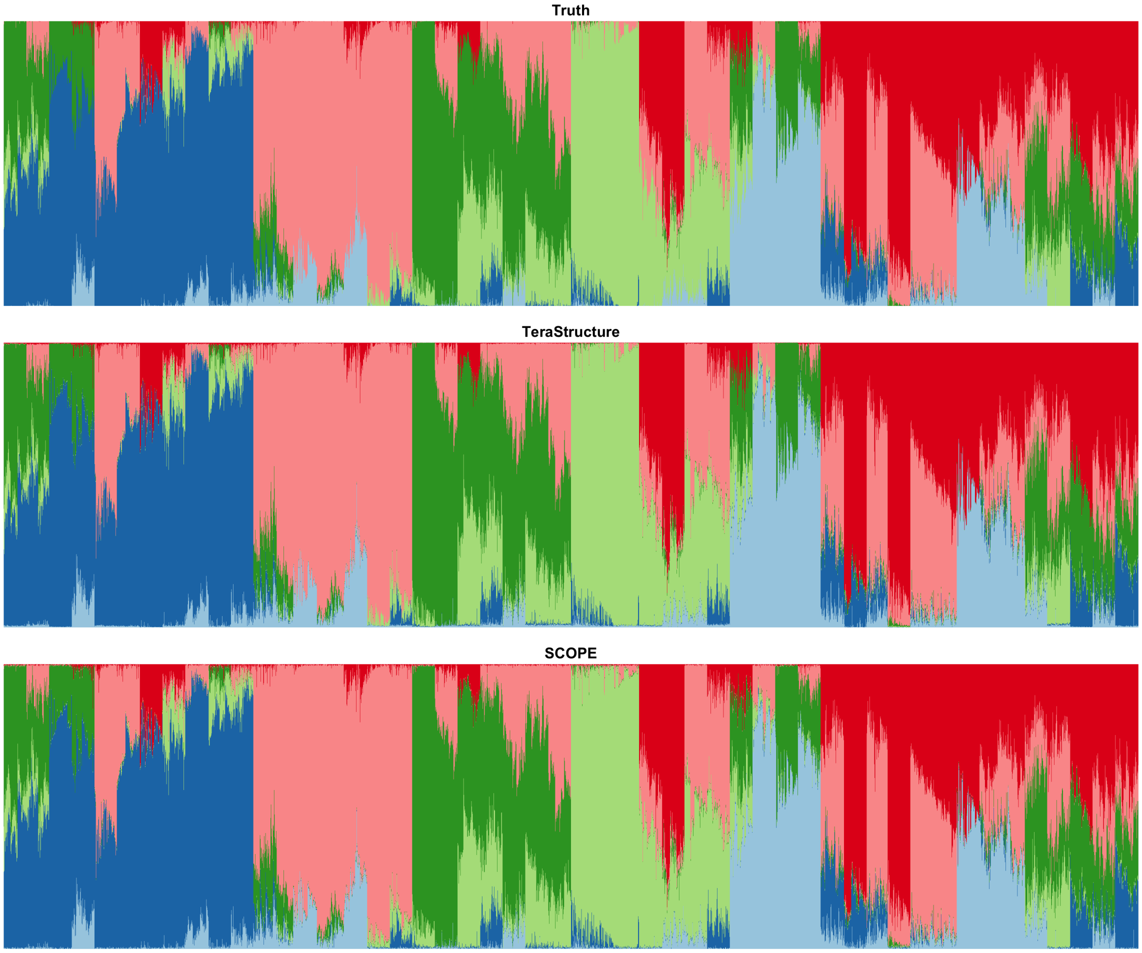


Figure S3: **Population structure inference for simulations under PSD model generated using 1000 Genomes Phase 3 data.** PSD model parameters were drawn from TGP data to generate a simulation dataset with 100,000 samples and 1 million SNPs. The true admixture proportions and resulting inferred admixture proportions from each method are shown. Colors and order of samples are matched between each method to the truth.

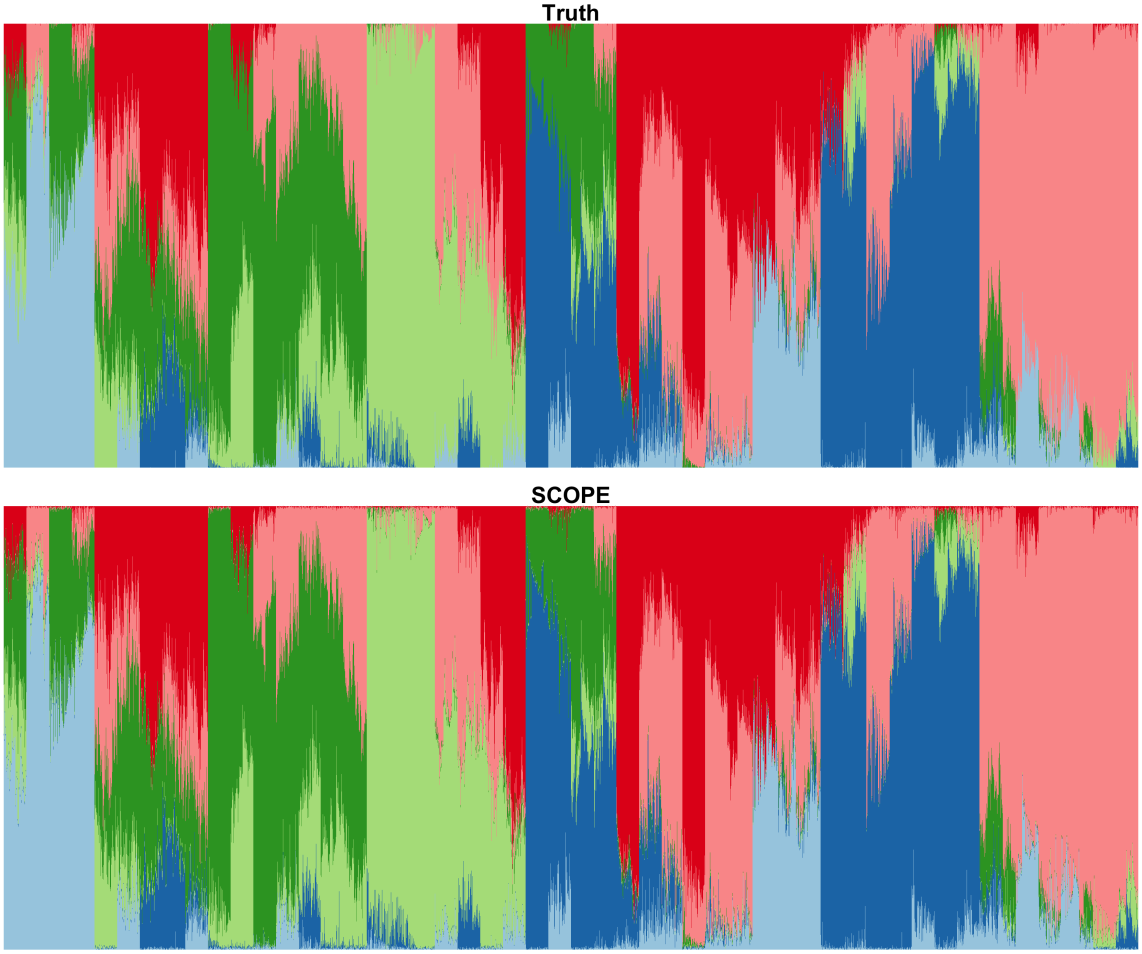


Figure S4: **Population structure inference for simulations under PSD model generated using 1000 Genomes Phase 3 data.** PSD model parameters were drawn from TGP data to generate a simulation dataset with 1 million samples and SNPs SNPs. The true admixture proportions and resulting inferred admixture proportions are shown. Colors and order of samples are matched between SCOPE and the true admixture proportions.

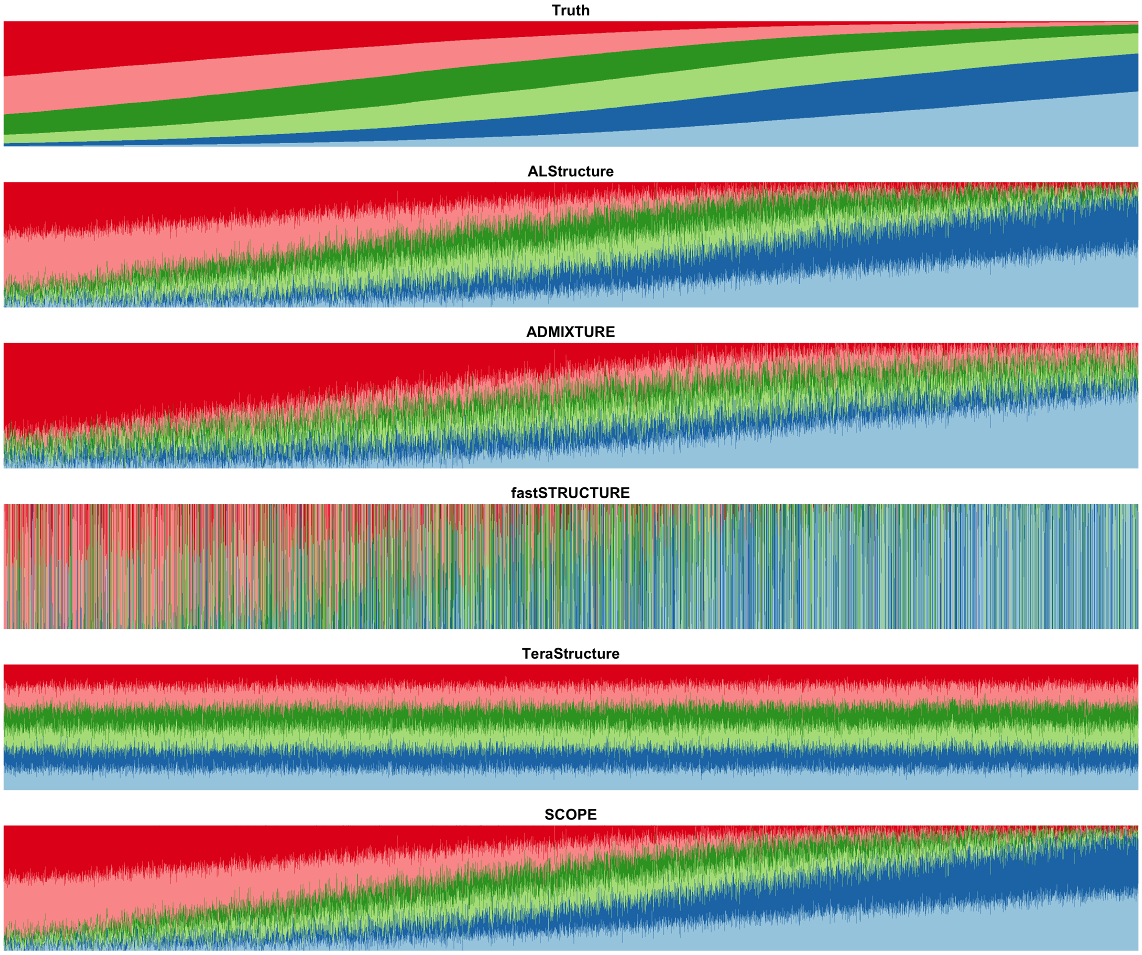


Figure S5: **Population structure inference for simulations under a spatial model generated using Human Genome Diversity Project data.** Model parameters were drawn from HGDP data to generate a simulation dataset with 10,000 samples and 10,000 SNPs under a spatial model (see Methods). The true admixture proportions and resulting inferred admixture proportions from each method are shown. Colors and order of samples are matched between each method to the truth.

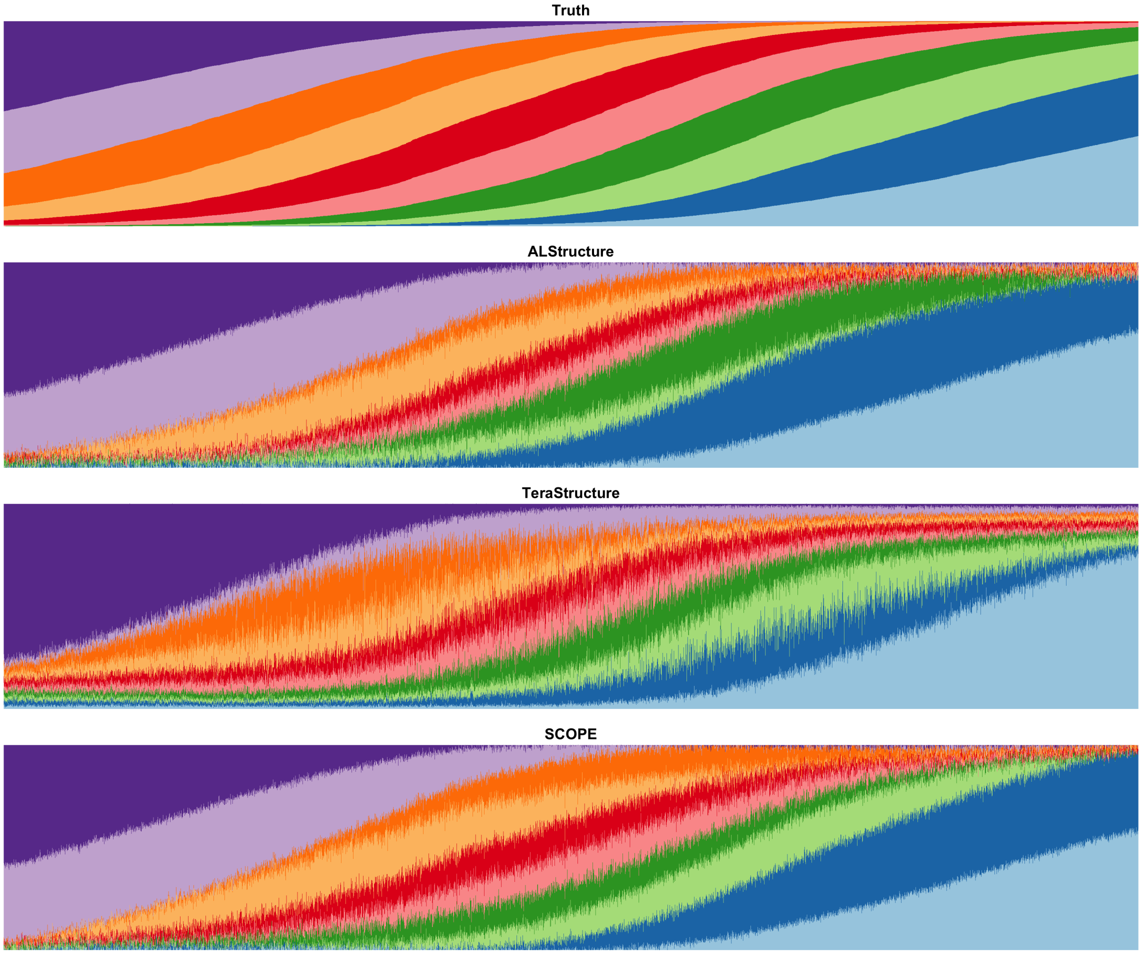


Figure S6: **Population structure inference for simulations under a spatial model generated using 1000 Genomes Phase 3 data.** Model parameters were drawn from TGP data to generate a simulation dataset with 10,000 samples and 100,000 SNPs under a spatial model (see Methods). The true admixture proportions and resulting inferred admixture proportions from each method are shown. Colors and order of samples are matched between each method to the truth.

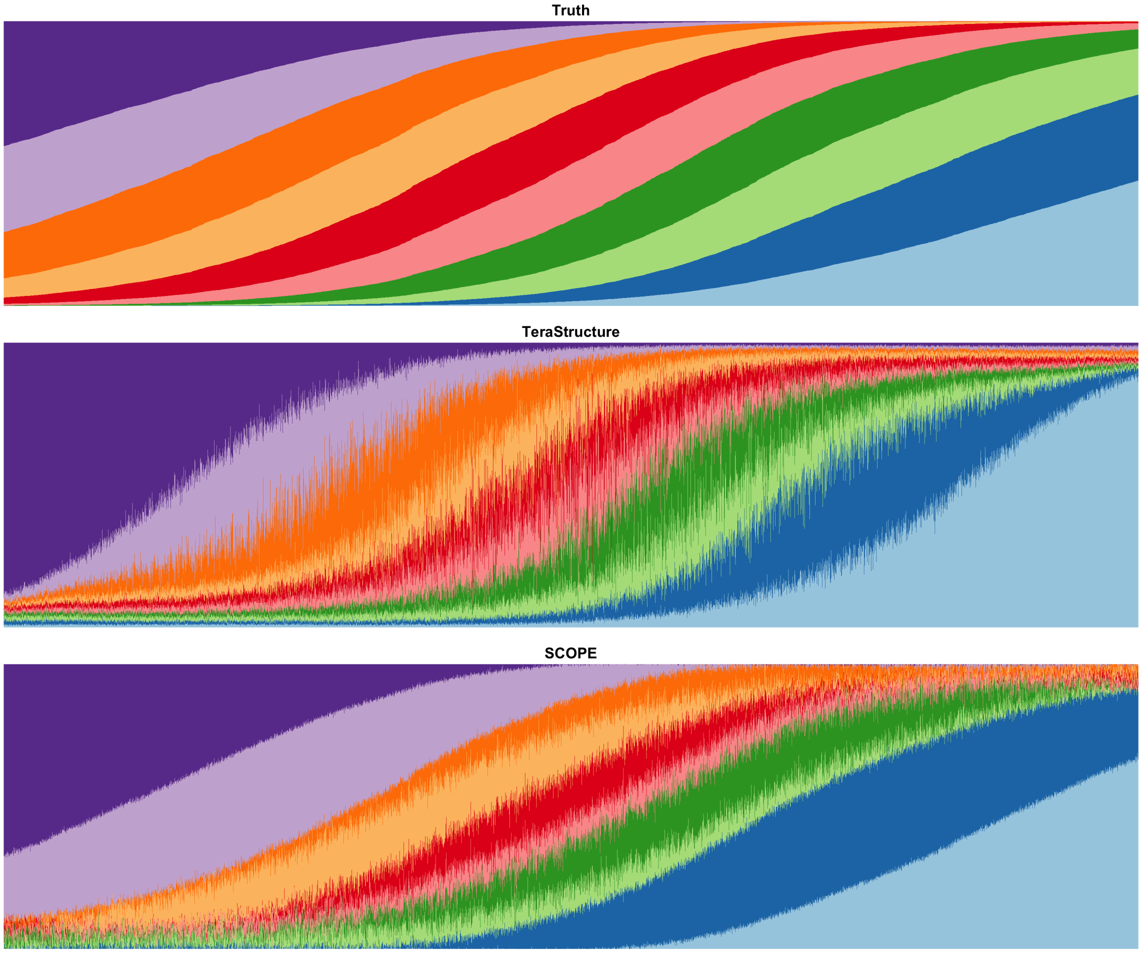


Figure S7: **Population structure inference for simulations under a spatial model generated using 1000 Genomes Phase 3 data.** Model parameters were drawn from TGP data to generate a simulation dataset with 10,000 samples and 1 millions SNPs under a spatial model (see Methods). The true admixture proportions and resulting inferred admixture proportions from each method are shown. Colors and order of samples are matched between each method to the truth.

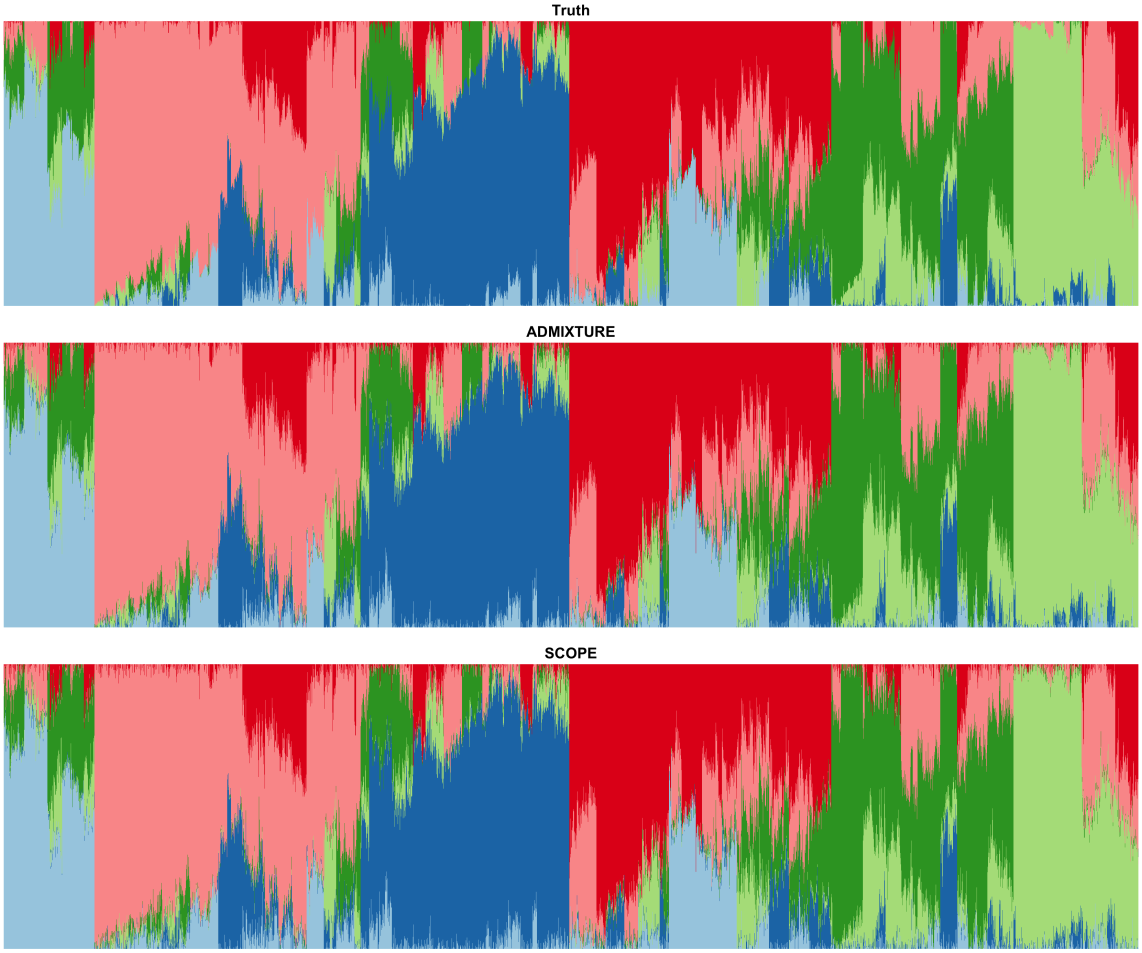


Figure S8: **Supervised population structure inference for simulations under the PSD model generated using 1000 Genomes Phase 3 data.** PSD model parameters were drawn from TGP data to generate a simulation dataset with 10,000 samples and 10,000 SNPs. Both were methods provided the true population allele frequencies as input. Colors and order of samples are matched between each method to the truth.

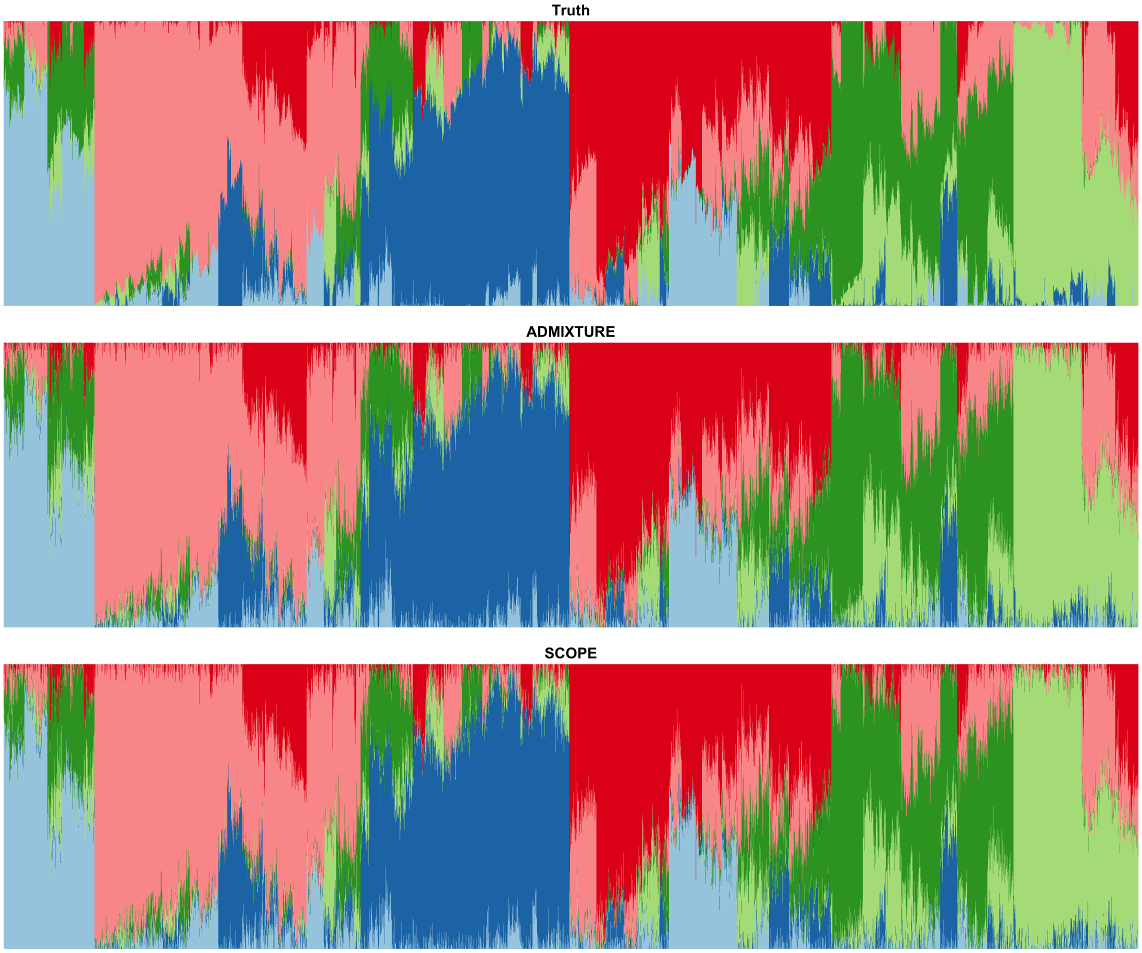


Figure S9: **Supervised population structure inference for simulations under the PSD model generated using Human Genome Diversity data.** PSD model parameters were drawn from HGDP data to generate a simulation dataset with 10,000 samples and 10,000 SNPs. Both were methods provided the true population allele frequencies as input. Colors and order of samples are matched between each method to the truth.

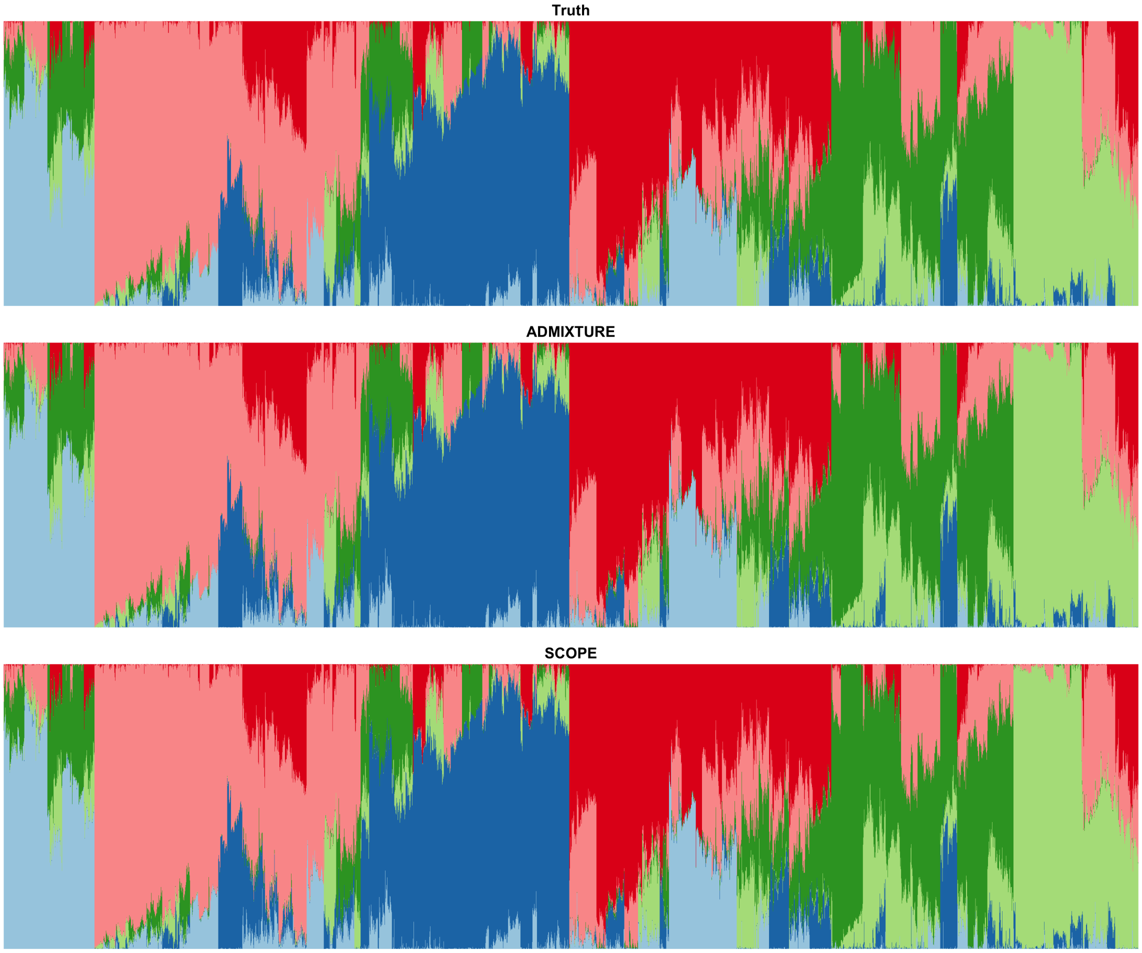


Figure S10: **Supervised population structure inference for simulations under the PSD model generated using 1000 Genomes Phase 3 data.** PSD model parameters were drawn from TGP data to generate a simulation dataset with 10,000 samples and 1 million SNPs. Both were methods provided the true population allele frequencies as input. Colors and order of samples are matched between each method to the truth.

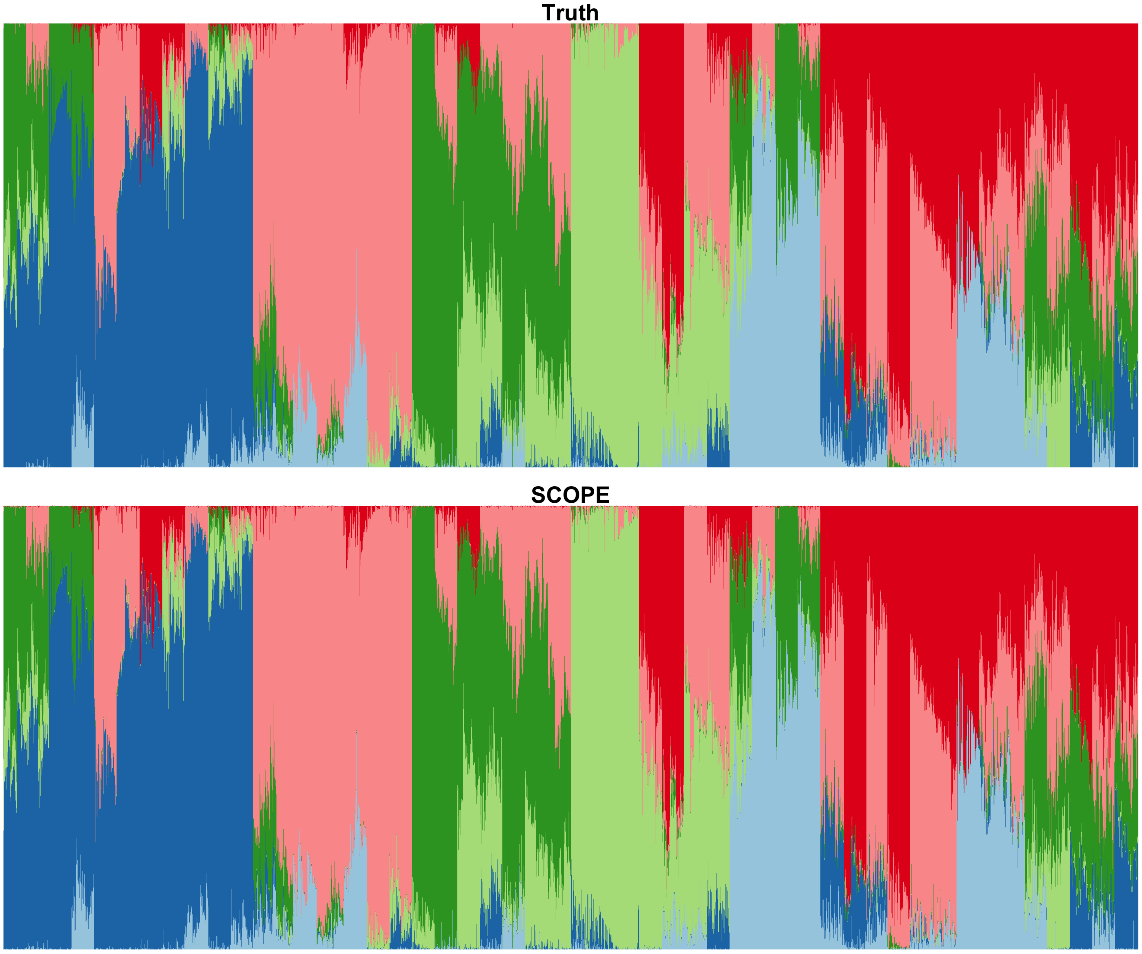


Figure S11: **Supervised population structure inference for simulations under the PSD model generated using 1000 Genomes Phase 3 data.** PSD model parameters were drawn from TGP data to generate a simulation dataset with 100,000 samples and 1 million SNPs. Both were methods provided the true population allele frequencies as input. Colors and order of samples are matched between each method to the truth.

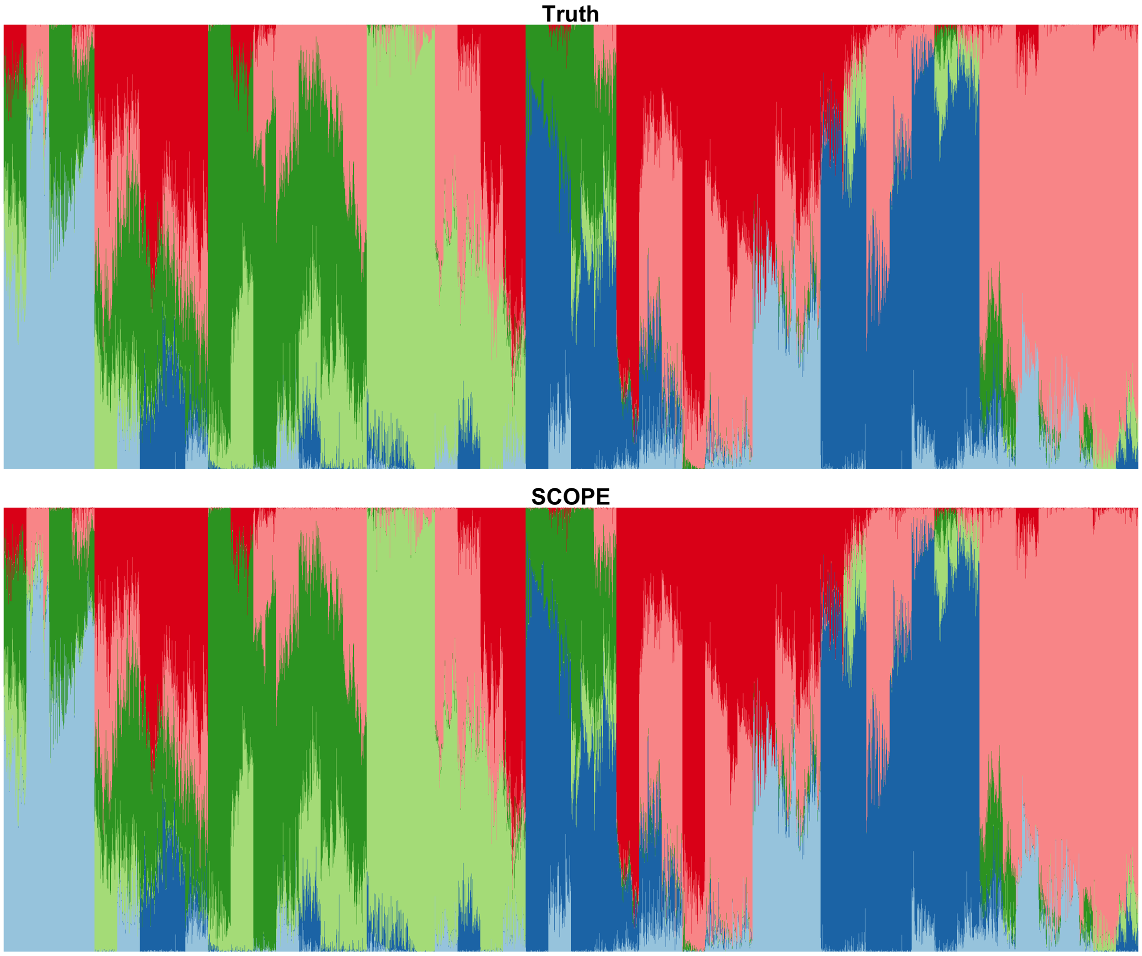


Figure S12: **Supervised population structure inference for simulations under the PSD model generated using 1000 Genomes Phase 3 data.** PSD model parameters were drawn from TGP data to generate a simulation dataset with 1 million individuals SNPs. SCOPE was provided the true population allele frequencies as input. Colors and order of samples are matched between SCOPE and the truth.

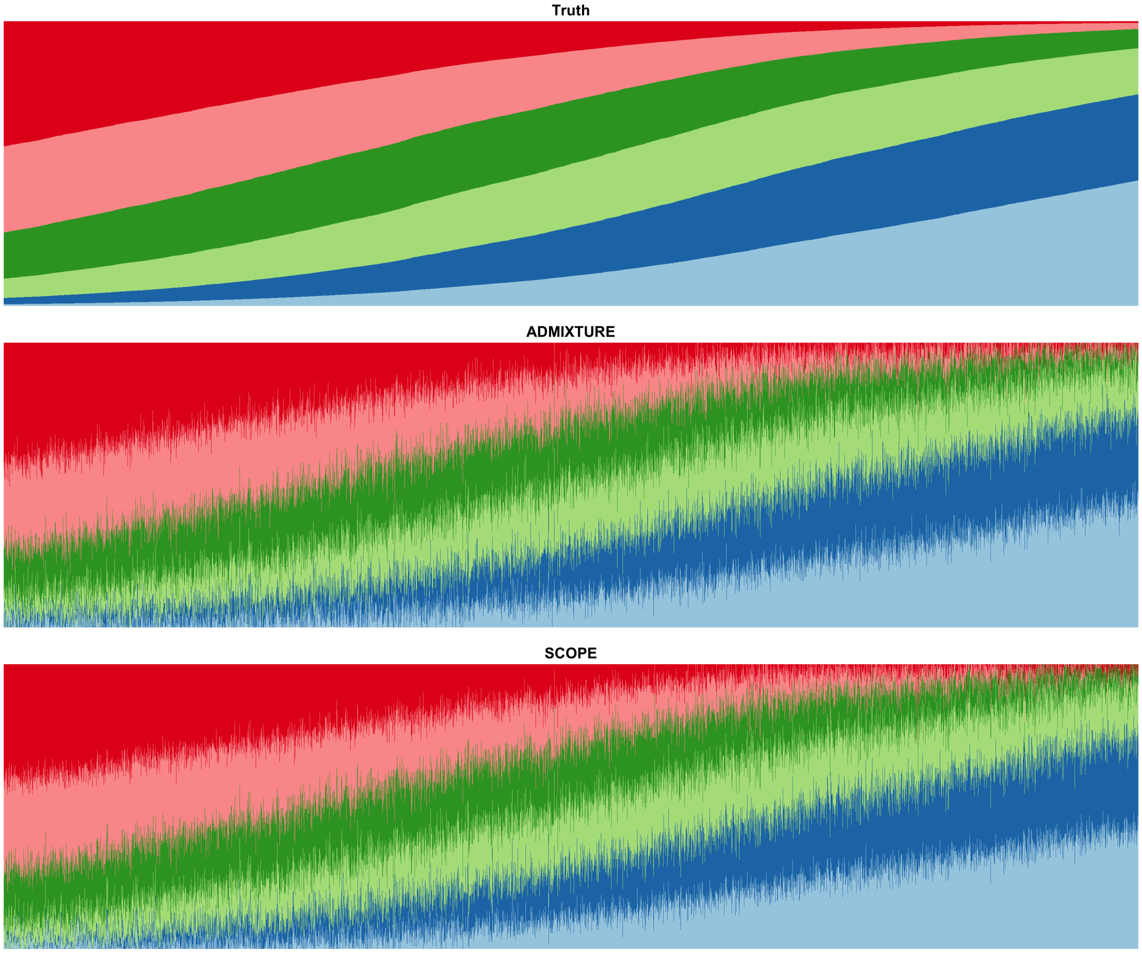


Figure S13: **Supervised population structure inference for simulations under a spatial model generated using Human Genome Diversity Project data.** Model parameters were drawn from HGDP data to generate a simulation dataset with 10,000 samples and 10,000 SNPs under a spatial model. Both methods were provided the true population allele frequencies as input. Colors and order of samples are matched between each method to the truth.

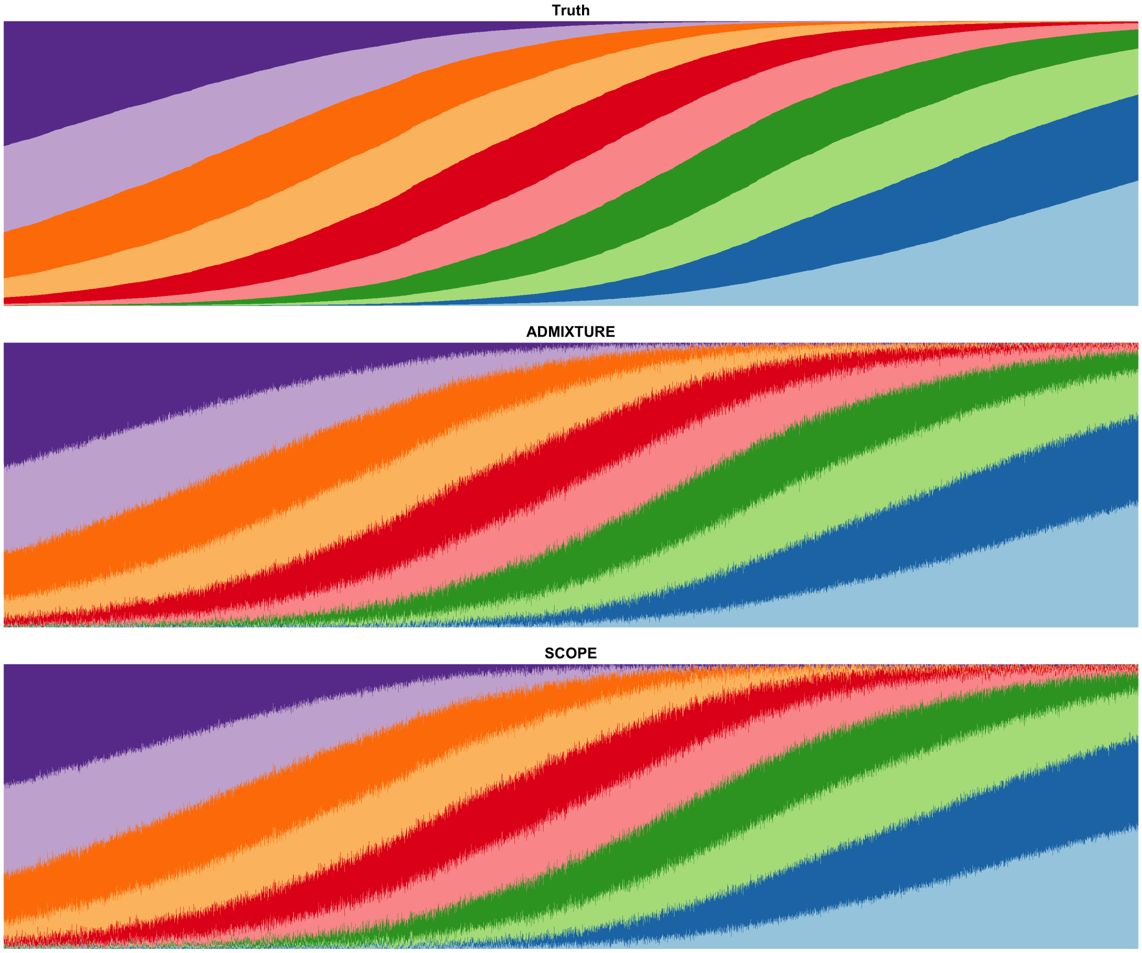


Figure S14: **Supervised population structure inference for simulations under a spatial model generated using 1000 Genomes Phase 3 data.** Model parameters were drawn from TGP data to generate a simulation dataset with 10,000 samples and 100,000 SNPs under a spatial model. Both methods were provided the true population allele frequencies as input. Colors and order of samples are matched between each method to the truth.

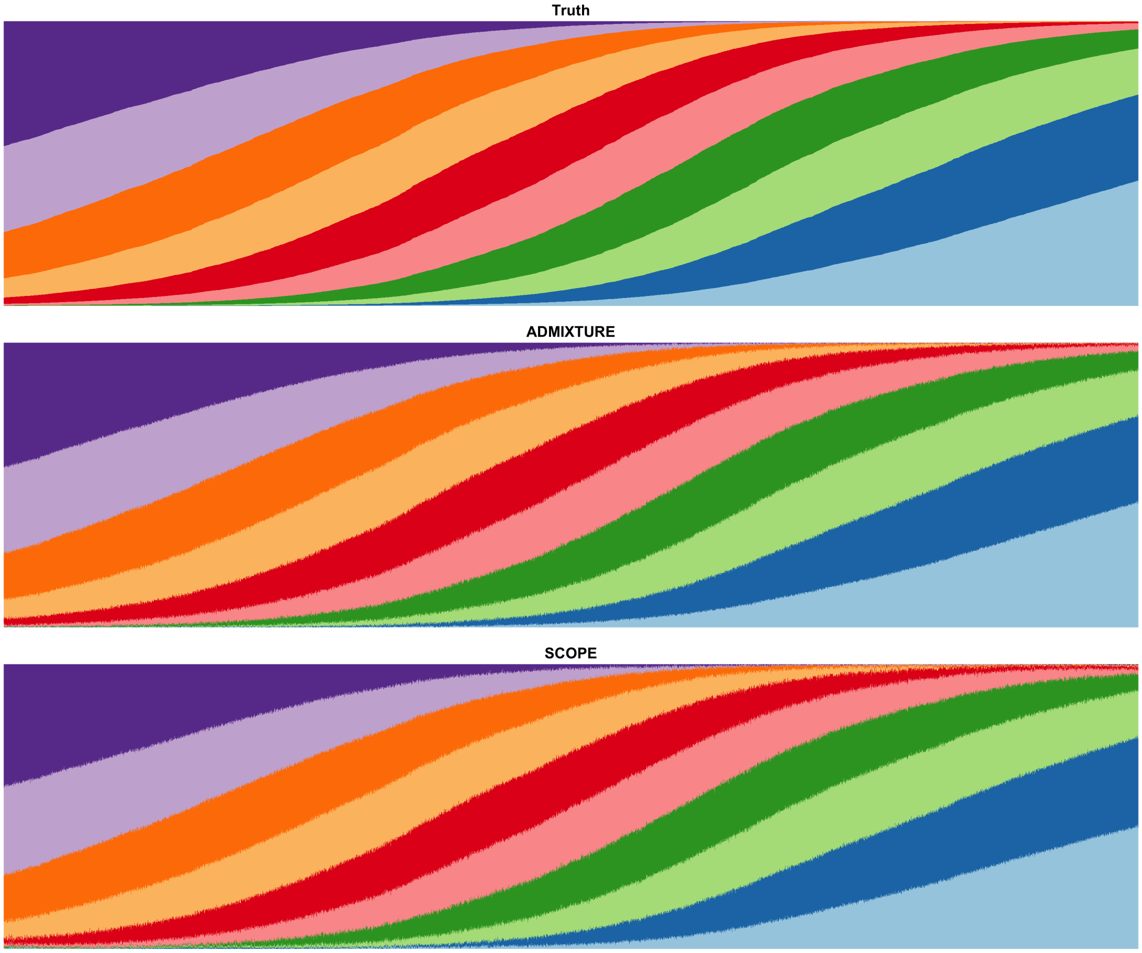


Figure S15: **Supervised population structure inference for simulations under a spatial model generated using 1000 Genomes Phase 3 data.** Model parameters were drawn from TGP data to generate a simulation dataset with 10,000 samples and 1 million SNPs under a spatial model. Both methods were provided the true population allele frequencies as input. Colors and order of samples are matched between each method to the truth.

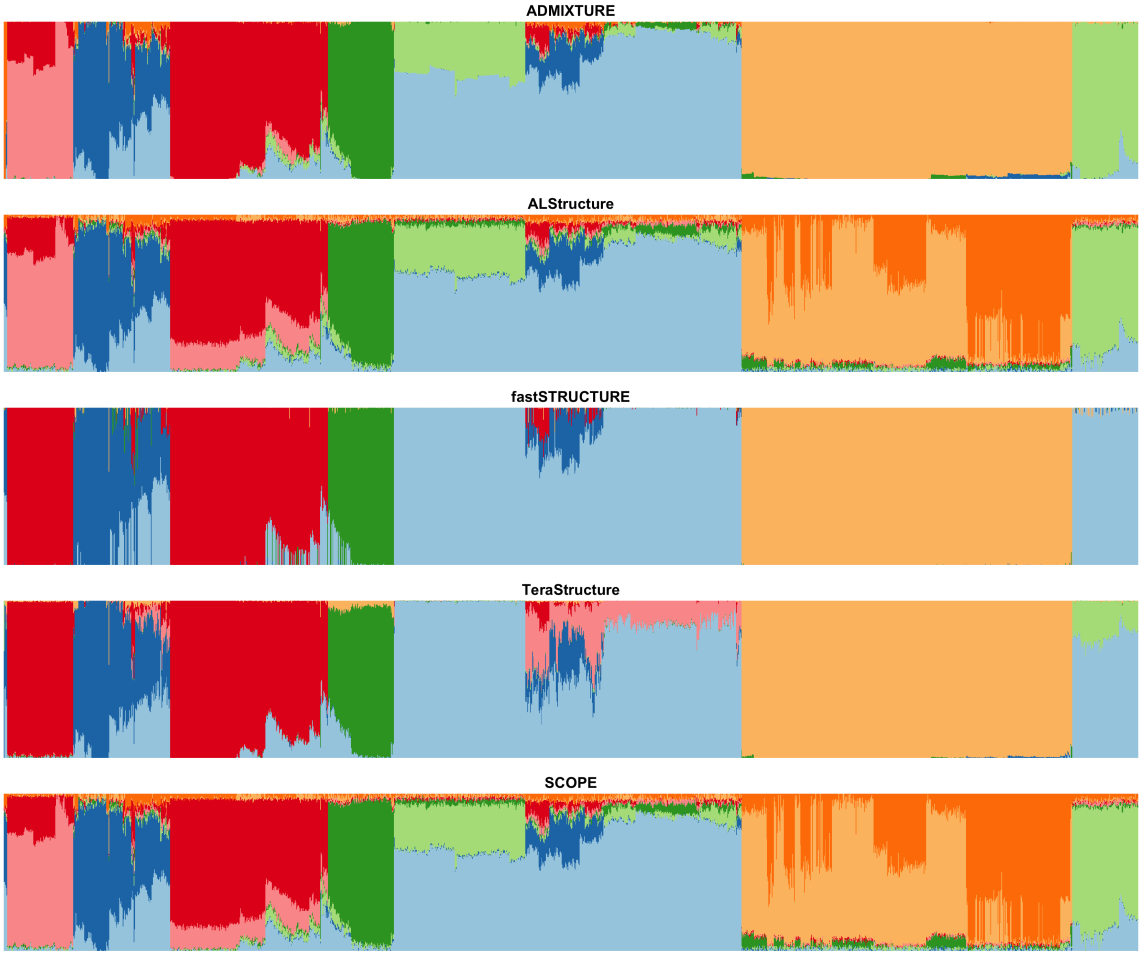


Figure S16: **Population structure inference of 1000 Genomes Phase 3 data using 8 latent populations.** Colors and order of samples are matched between each method and ADMIXTURE. ADMIXTURE was ordered through hierarchical clustering (see Methods).

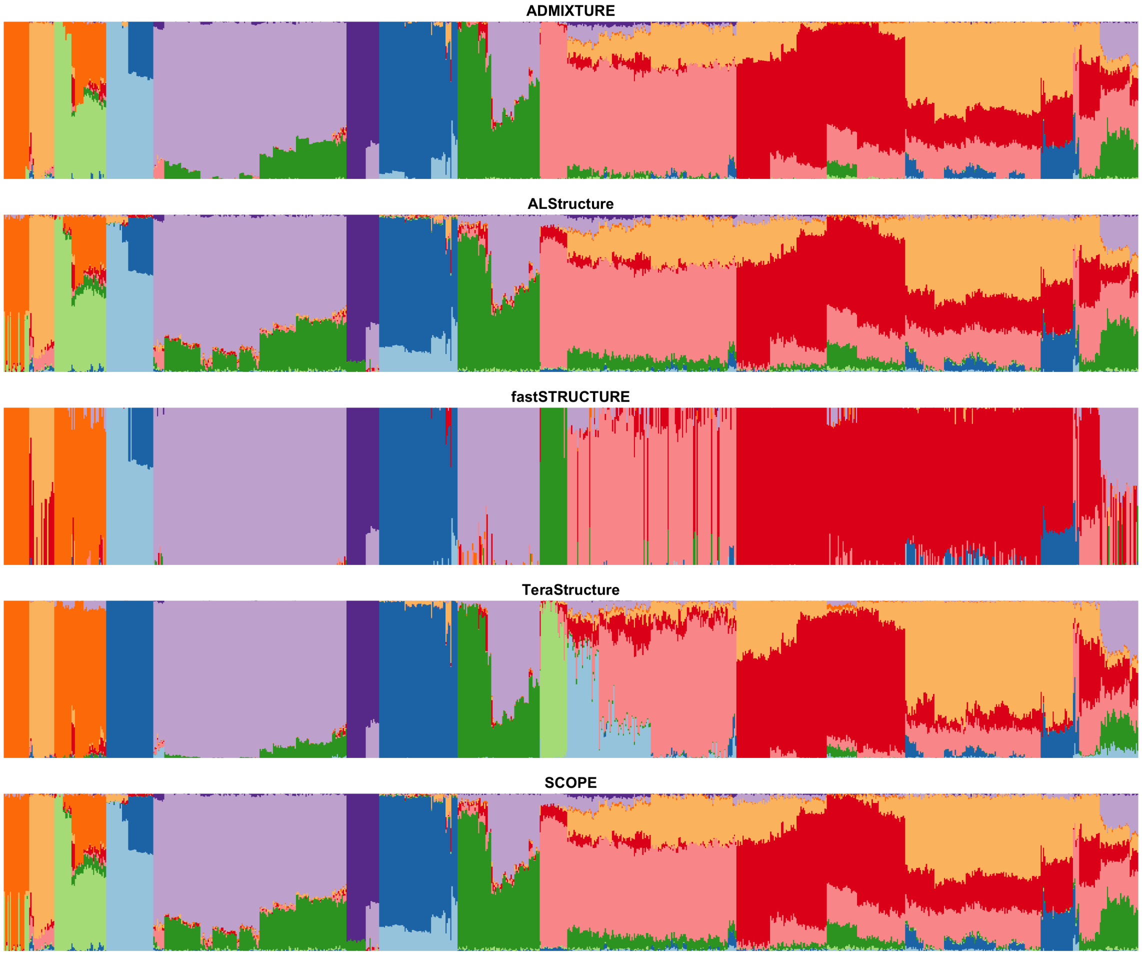


Figure S17: **Population structure inference of Human Genomes Diversity Population data using 10 latent populations.** Colors and order of samples are matched between each method and ADMIXTURE. ADMIXTURE was ordered through hierarchical clustering (see Methods).

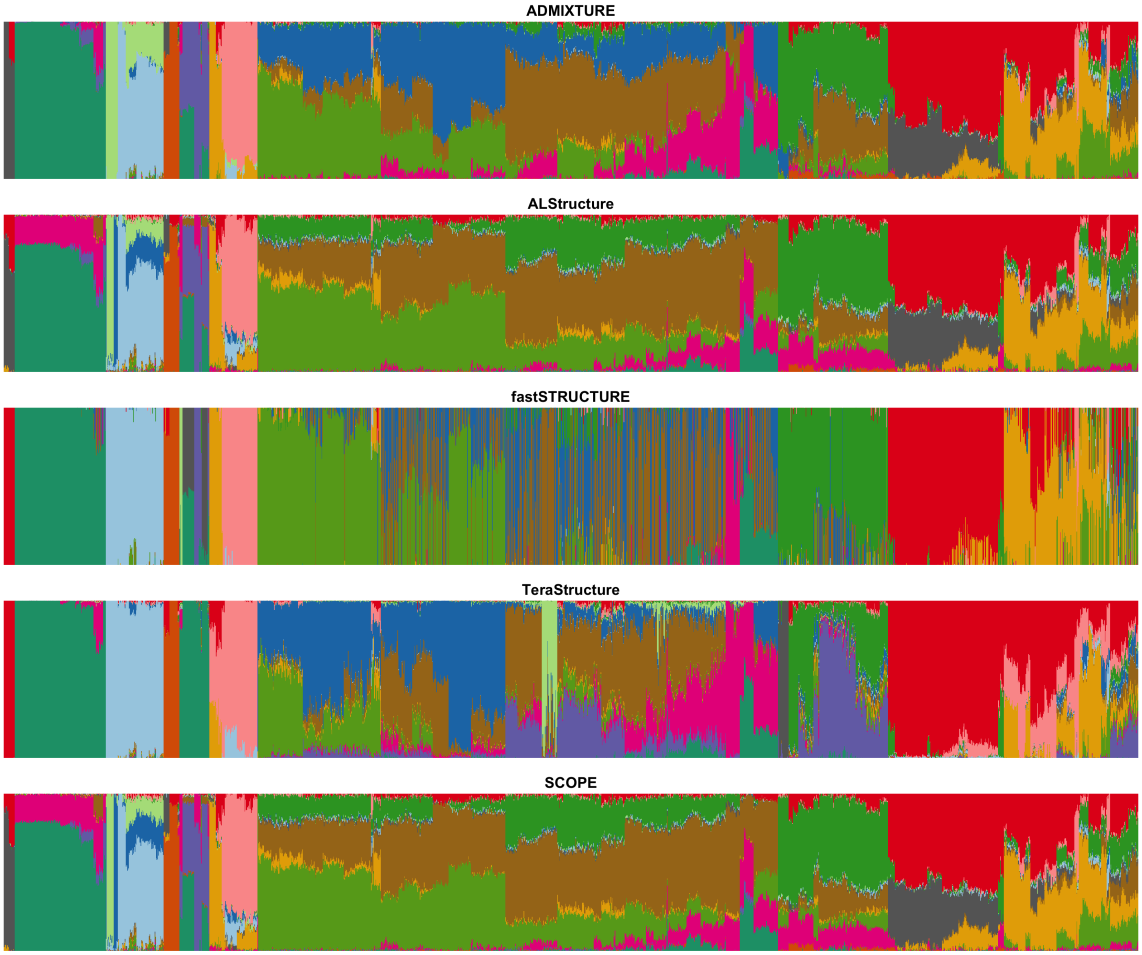


Figure S18: **Population structure inference of Human Origins data using 14 latent populations.** Colors and order of samples are matched between each method and ADMIXTURE. ADMIXTURE was ordered through hierarchical clustering (see Methods).

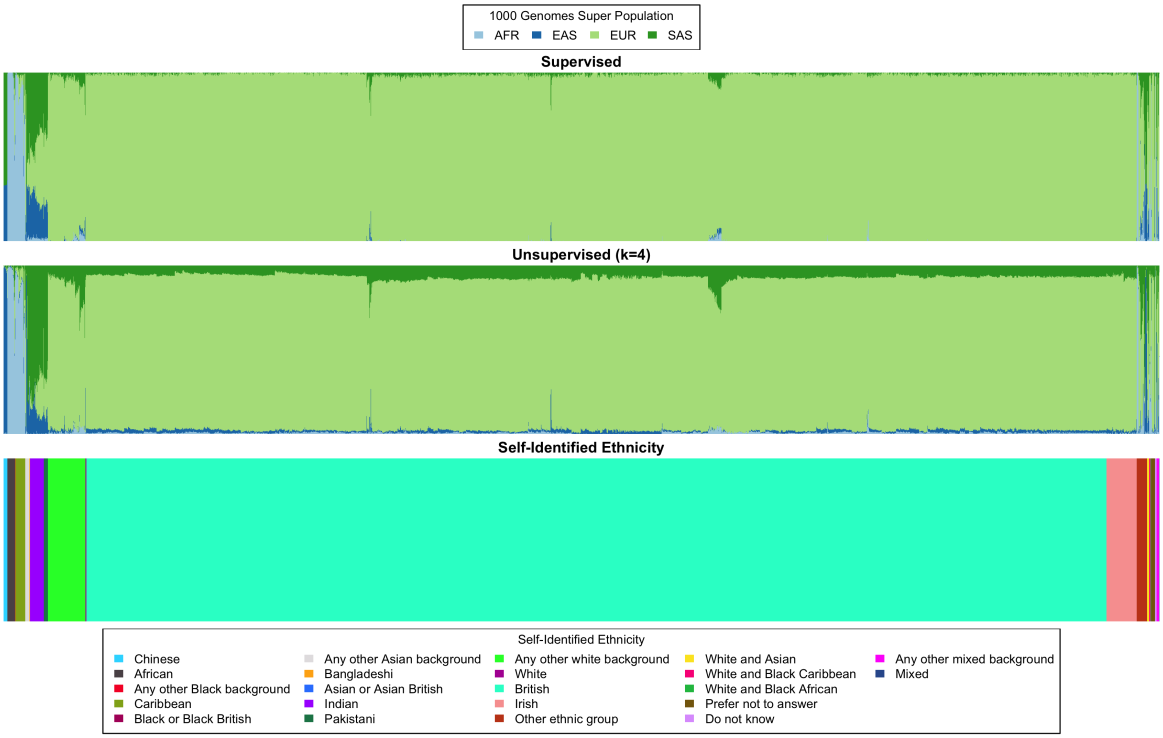


Figure S19: **Population structure inference on the UK Biobank with all individuals.** We ran population structure inference using SCOPE in both supervised mode using 1000 Genomes Phase 3 allele frequencies (top) and unsupervised with 4 latent populations (middle). For reference, we plot the self-identified race/ethnicity (bottom). Colors and order of samples are matched between each row of the figure. This is an extended version of Figure 4 that includes all self-identified British samples.

Table S1: **Memory usage of methods on simulated and real datasets.** ADMIXTURE, TeraStructure, and SCOPE were run using 8 threads. ALStructure and fastStructure were run on a single thread due to their lack of multithreading implementations. TeraStructure's '-rfreq' parameter was set to 10% of the number of SNPs. A '-' denotes that the method was not run due to projected time or memory usage. Default parameters were used otherwise. Memory is displayed in gigabytes (GB). Bold values denote the best value for each dataset.

| Dataset Type | Base Dataset | k | n | m | ADMIXTURE | fastStructure | TeraStructure | ALStructure | SCOPE |
|---|---|---|---|---|---|---|---|---|---|
| PSD | HGDP | 6 | 10,000 | 10,000 | **0.12** | 0.17 | **0.12** | 7.30 | 0.14 |
| PSD | TGP | 6 | 10,000 | 10,000 | **0.12** | 0.16 | **0.12** | 7.30 | 0.14 |
| PSD | TGP | 6 | 10,000 | 1,000,000 | 10.66 | 10.66 | **9.96** | - | 12.60 |
| PSD | TGP | 6 | 100,000 | 1,000,000 | - | - | 94.38 | - | **93.47** |
| PSD | TGP | 6 | 1,000,000 | 1,000,000 | - | - | - | - | **746.19** |
| Spatial | HGDP | 6 | 10,000 | 10,000 | **0.12** | 0.17 | **0.12** | 7.30 | 0.14 |
| Spatial | TGP | 6 | 10,000 | 10,000 | **0.12** | 0.16 | **0.12** | 7.30 | 0.14 |
| Spatial | TGP | 10 | 10,000 | 100,000 | 1.17 | 1.33 | **1.05** | 33.20 | 1.28 |
| Spatial | TGP | 10 | 10,000 | 1,000,000 | - | - | **10.30** | - | 12.69 |
| Real | HGDP | 10 | 940 | 642,951 | 1.94 | 1.99 | **1.17** | 24.38 | 1.30 |
| Real | HO | 14 | 1,931 | 385,089 | 1.83 | 1.89 | **1.21** | 27.45 | 1.53 |
| Real | TGP | 8 | 1,718 | 1,854,622 | 6.20 | 6.18 | **4.44** | 145.49 | 6.34 |
| Real | UKB | 4 | 488,363 | 569,346 | - | - | - | - | **230.57** |

Table S2: **Accuracy of supervised population structure inference for SCOPE and ADMIXTURE using supplied allele frequencies on simulations.** True allele frequencies were supplied to each method. Root-mean-square error (RMSE) and Jensen-Shannon Divergence (JSD) were computed against the true admixture proportions. Estimated proportions of 0 were set to $1 \times 10^{-9}$ for JSD calculations (see Methods). A "-" denotes that the method was not run for that dataset due to time or memory constraints. Values are displayed as percentages. Bold values denote the best value for each dataset.

| | | | | | SCOPE | | ADMIXTURE | |
|---|---|---|---|---|---|---|---|---|
| Dataset Type | Base Dataset | k | n | m | RMSE | JSD | RMSE | JSD |
| PSD | HGDP | 6 | 10,000 | 10,000 | 2.9 | 2.1 | **2.6** | **1.8** |
| PSD | TGP | 6 | 10,000 | 10,000 | 2.0 | 1.3 | **1.6** | **0.8** |
| PSD | TGP | 6 | 10,000 | 1,000,000 | **0.2** | 0.1 | **0.2** | **0.04** |
| PSD | TGP | 6 | 100,000 | 1,000,000 | **0.2** | 0.1 | **0.2** | **0.04** |
| PSD | TGP | 6 | 1,000,000 | 1,000,000 | **0.2** | **0.1** | - | - |
| Spatial | HGDP | 6 | 10,000 | 10,000 | **2.4** | **0.8** | 3.2 | 1.3 |
| Spatial | TGP | 6 | 10,000 | 10,000 | **1.7** | **0.5** | 2.2 | 0.6 |
| Spatial | TGP | 10 | 10,000 | 100,000 | **0.6** | **0.4** | 0.7 | **0.4** |
| Spatial | TGP | 10 | 10,000 | 1,000,000 | 0.3 | **0.1** | **0.2** | **0.1** |

Table S3: **Prediction accuracy of self-identified race and ethnicity using inferred admixture proportions.** We trained multinomial logistic regression models using the inferred admixture proportions from each method to predict SIRE labels. For TGP, we predicted 5 superpopulation labels corresponding to continental ancestry from 8 inferred latent populations. For HGDP, we predicted 7 continental ancestry populations from 10 inferred latent populations. Training accuracy as a percentage is reported.

| Method | TGP | HGDP |
|---|---|---|
| ADMIXTURE | 100 | 46.4 |
| ALStructure | 100 | 47.6 |
| fastStructure | 99.4 | 41.8 |
| TeraStructure | 100 | 47.8 |
| SCOPE | 100 | 47.2 |
